# 3D chromatin organization changes modulate adipogenesis and osteogenesis

**DOI:** 10.1101/2020.05.25.114272

**Authors:** Ruo-Han Hao, Yan Guo, Jing Guo, Yu Rong, Shi Yao, Yi-Xiao Chen, Shan-Shan Dong, Dong-Li Zhu, Hao Chen, Tie-Lin Yang

## Abstract

Human mesenchymal stem cells (hMSCs) can be differentiated into adipocytes and osteoblasts. While the transcriptomic and epigenomic changes during adipogenesis and osteogenesis have been characterized, what happens to the chromatin loops is hardly known. Here we induced hMSCs to adipogenic and osteogenic differentiation, and performed 2 kb resolution Hi-C experiments for loop detection and generated RNA-seq, histone modification ChIP-seq and ATAC-seq data for integrative analysis before and after differentiation. We quantitatively identified differential contact loops and unique loops. After integrating with multi-omics data, we demonstrate that strengthened loops after differentiation are associated with gene expression activation. Specially, unique loops are linked with cell fate determination. We also proposed loop-mediated regulatory networks and identified *IRS2* and *RUNX2* as being activated by cell-specific loops to facilitate adipocytes and osteoblasts commitment, respectively. These results are expected to help better understand the long-range regulation in controlling hMSC differentiation, and provide novel targets for studying adipocytes and osteoblasts determination.

## Introduction

Human mesenchymal stem cells (hMSCs) are multipotential cells and capable of differentiating into a number of common lineages, including adipocytes and osteoblasts. Previous studies have characterized many key factors that manipulate hMSC differentiation. For example, the transcriptome profiling throughout the lineage commitment of MSC cell line into adipocytes^1, 2^ and osteoblasts^3, 4^ has already been reported, and a bunch of signature genes have been identified in these processes^5^. In particular, the investigation of core transcription factors (TFs) during hMSC differentiation has also made marked success, and uncovered several master regulators, like peroxisome proliferator-activated receptor γ (PPARγ)^6, 7, 8^, CCAAT-enhancer-binding proteins α (C/EBPα)^9, 10^ for adipogenesis, and runt-related transcription factor 2 (RUNX2)^11^, Osterix (OSX/SP7)^12^ for osteogenesis. Besides, epigenomic programming provides another view of dynamic histone modifications and enhancer activity during mouse MSCs differentiation^13, 14^. Recently, studies focusing on the open chromatin regions have attracted attention to the rewiring of chromatin structure during adipogenesis and osteogenesis^15^. Even though great efforts have been made to reveal the biological process during lineage commitment of MSCs into osteoblasts and adipocytes, these studies were carried out with different cellular models having divergent genetic backgrounds. Thus, a holistic insight is needed through conducting those assays with an uniform model.

Taking advantage of the Hi-C technology, the spatial organization of the human genome has been revealed at different resolutions in different conditions. Although previous studies have reported the close relationship between the topologically associated domains (TADs) and many biological processes^16, 17, 18^, the structures of detailed chromatin interactions are still hidden due to the limited resolution. As the increasement of sequence depth, loop structures are able to be detected. Unlike TADs dividing chromosomal into different territories, chromatin loops directly bring distal elements into close proximity with their target promoters^19^, and the shortened distance between enhancer and promoter contributes to gene activation^19, 20, 21^. In addition, studies focusing on loop structures have uncovered the dynamic interaction changes resulting in cell function determination^22, 23^. In terms of cell fate commitment, several studies have reported the chromatin organization rewiring during stem cell differentiation^16, 23^. However, to date, the investigation of how the hMSC chromatin structure response to adipogenic and osteogenic induction is still blank, especially for chromatin loops, yet is required to understand the underlying differentiation mechanism.

Here we performed high resolution Hi-C experiments before and after hMSC was differentiated into adipocytes and osteoblasts, and included RNA-seq, ChIP-seq and ATAC-seq at each stage simultaneously in order to provide a comprehensive insight of loop-mediated regulation patterns using data come from the same cellular model. We found differential contacted loops in each cell, and showed that the strengthened loops in both differentiation terminal cells are associated with gene activation, differential enhancer reprograming and active TF binding. In particular, unique loops are essential for cell fate determination. Eventually, by constructing loop-mediated regulatory networks, we reveal the cell-specific regulation cascades and identify the controlling factors for adipogenesis and osteogenesis.

## Results

### 3D chromatin architectures of hMSC, adipocytes and osteoblasts

In order to study chromatin conformation changes after hMSC differentiation, we carried out high-resolution Hi-C experiments before and after inducing hMSC differentiation into adipocytes (AC) and osteoblasts (OB) separately (Fig. 1a and Supplementary Fig. 1). At least six replicates were generated for each cell type. Hi-C data of each cell were combined to produce an average of 3.6 billion qualified paired-end reads after filtering out potentially artificial reads. We implemented HiCUP^24^ to process Hi-C data, which resulted in ~2.5 billion valid read pairs (Supplementary Table 1). We estimated the average intrachromosomal contact probability in each cell. The contact probability curves were similar across cells (Supplementary Fig. 2a) and consistent with a previous report in lymphoblastoid cell line^25^, suggesting that the Hi-C data were qualified to detect intrachromosomal interactions. We further verified the reproducibility through the high correlation of normalized contact frequency among replicates (Supplementary Fig. 2b).

**Fig. 1.**
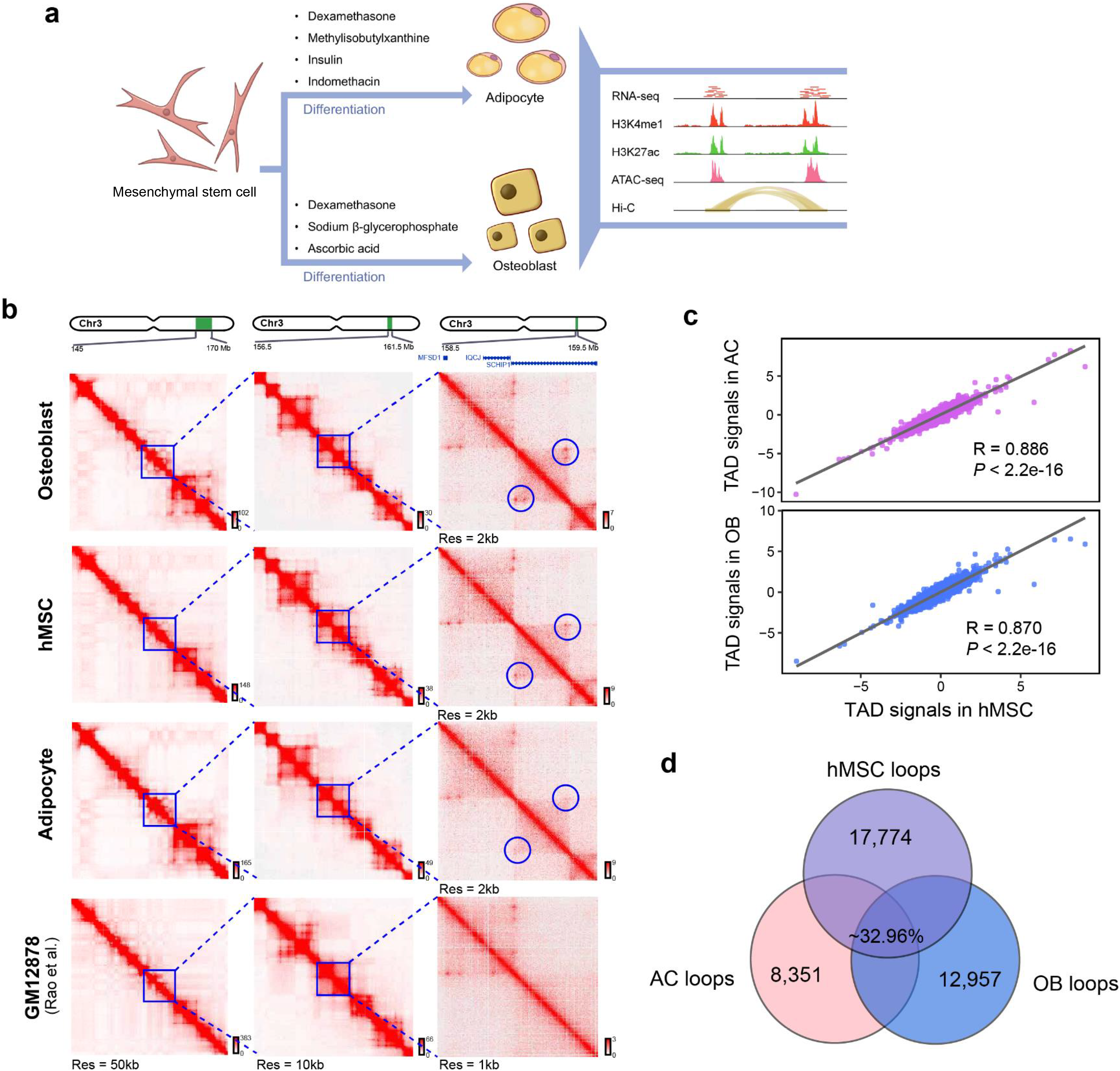
Chromatin conformation features of hMSC and differentiated adipocytes and osteoblasts. **a** hMSCs were differentiated into adipocytes and osteoblasts by supplying with specific differentiation media. The collections of cells were subjected to Hi-C, RNA-seq, ChIP-seq and ATAC-seq measurement. **b** Normalized Hi-C contact heatmap at 50 kb, 10 kb and 2 kb/1 kb resolutions for different cells. An expected cell-specific interaction was circled, which also shows cell-lineage specificity. **c** The correlation of TAD signals between hMSC and differentiated adipocytes (AC) and osteoblasts (OB) cells, respectively. *P* values and correlation coefficients were estimated by Pearson correlation test. **d** Number of shared and cell-specific loops identified in 3 cells.

Using the “map resolution” definition described by Rao *et al.*^26^, we constructed chromatin contact matrices at 50 kb, 10 kb and 2 kb resolutions, respectively. As shown in Fig. 1b, genome organization details showed up after zooming in the maps to higher resolutions. Particularly, local chromatin interactions were able to be detected at 2 kb resolution (the dark “pixels points” shown in Fig. 1b, rightmost).

The genome is partitioned into “megadomains”, of which the chromatin status can be indicated by A and B compartments^26^. The “megadomains” are further partitioned into small topologically associating domains (TADs) of condensed chromatin. To find out the chromatin conformation changes after hMSC differentiation from different genome scales, we firstly conducted PCA analysis and directional index method^17^ to identify A/B compartment and TADs, respectively. We observed A/B compartment switch in response to differentiation induction (Supplementary Fig. 2c), reflecting different gene expression activity^16, 25^. As an example of adipogenesis associated gene *PPARG*, this gene is marked with B compartment (inactive status with negative PC1) in hMSC and OB but switched to active A compartment (positive PC1) in AC (Supplementary Fig. 2d). For TAD calling, among the 2,854 and 4,968 TADs identified in AC and OB (Supplementary Table 2), 88% and 70% were overlapped with TADs found in hMSC. Besides, the correlation of TAD signals showed high consistency across cells (Fig. 1c and Supplementary Fig. 2e). The conserved TADs between cells has been proved^17^, and our results confirmed this observation especially between hMSC and differentiated adipocytes and osteoblasts.

We next identified loop structures, and to also get a sense of how long-range interactome alters, we called chromatin interactions under 2 kb resolution (see Methods). We found 21,738, 12,460 and 16,930 loops in hMSC, AC and OB, respectively (Supplementary Table 3), and identified ~0.5 M significant interactions with 5% false discovery rate (FDR) control. In contrast with the stabilization of TADs, we uncovered many cell-specific loops and interactions which are difficult to be observed under low resolutions. At least 65% of loops were distinct, and ~33% were shared across cells (Fig. 1d). As an loop shown in Fig. 1b (rightmost column), when comparing with hMSC, the contact frequency was elevated in OB but was weakened in AC. Notably, when we compared with 1 kb resolution map in GM12878, it turned out to be a cell-linage-specific interaction. These genome intervals harbor a *IQCJ-SCHIP1* readthrough gene. *IQCJ* and *SCHIP1* mutated mouse exhibit skeletal defects, such as decreased bone mineral density and abnormal skeleton morphology^27^, highlighting the potential association between chromatin conformation alteration and cell function.

### Active regulatory elements are enriched in loop anchors

Further looking into chromatin loops and interactions, we were firstly interested to know what biological process was likely to take place at loop anchors or interacting fragments. We obtained chromatin states of enhancer, promoter, transcription regulatory and quiescent regions of hMSC, adipocyte and osteoblast from ChromHMM annotations to perform enrichment analysis (Supplementary Table 4; see Methods). Comparing with the genome, both loop anchors and interacting fragments were significantly enriched with active transcription regulatory elements and depleted of inactive elements across cells (Fig. 2a), supporting the opinion that promoter and regulatory elements are often interacting to facilitate gene expression^28^. We also analyzed region enrichment for both interacting fragments/anchors and their proximal length-match regions with 2 kb intervals (see Methods). Interestingly, comparing to the proximal regions, both loop anchors and interacting fragments tended to be more enriched with regulation signals but less occupied by transcription signals (Fig. 2b and Supplementary Fig. 3a; Supplementary Table 5). The difference was observed for CTCF peaks enrichment as well, which interacting fragments and loop anchors were more likely to harbor CTCF binding sites (Fig. 2c and Supplementary Fig. 3b). The significance was observed cross 3 cells and suggested that 3D chromatin folding is one of the important transcription regulatory features.

**Fig. 2.**
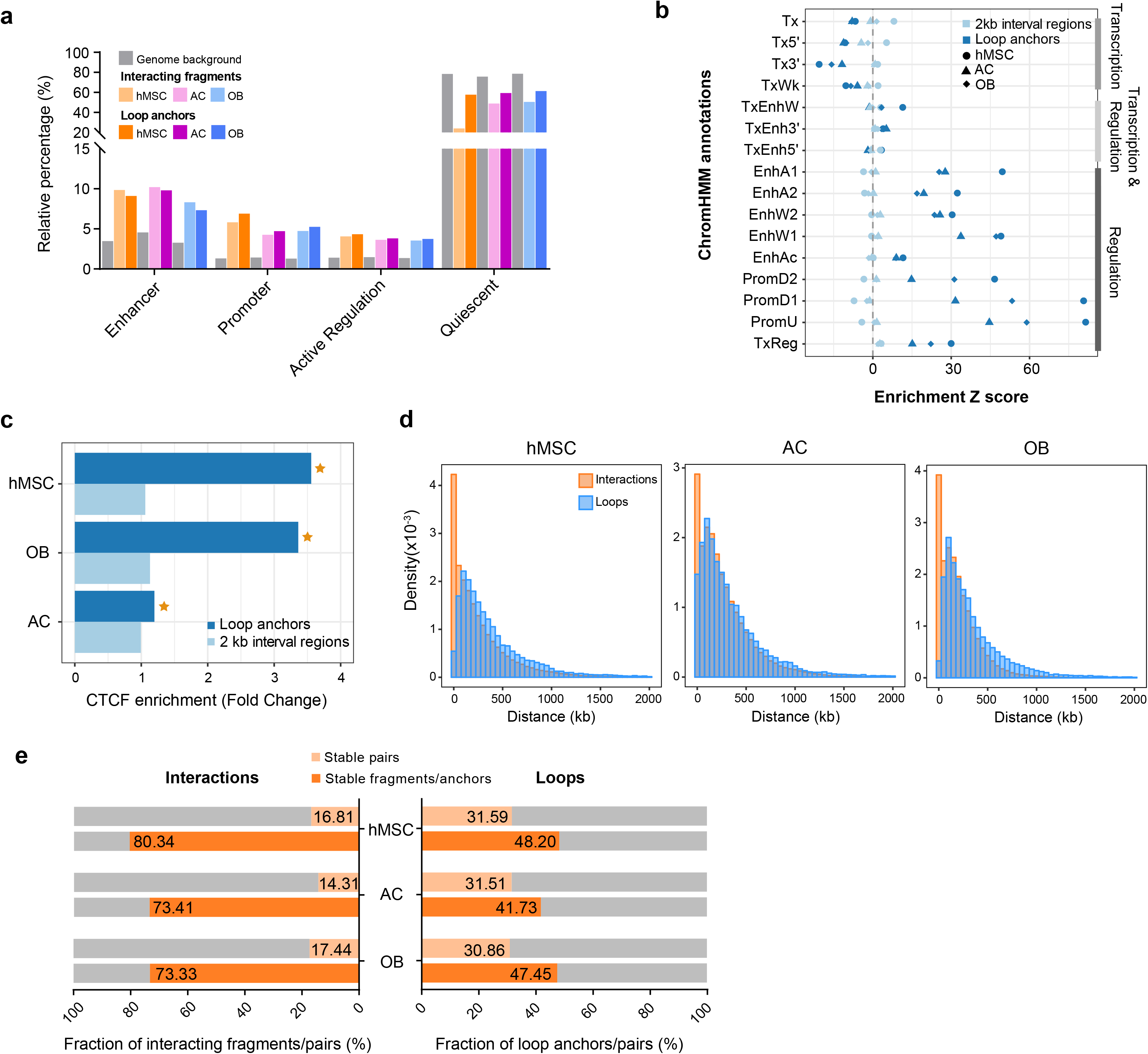
Chromatin loop anchors and interaction fragments are enriched with active regulation elements. **a** Regulatory elements annotation at loop anchors and interacting fragments. The relative fraction was compared with genomic background for enrichment analysis. Permutation test was executed to estimate enrichment significance. All comparisons have reached statistical significance (*P* < 0.05). **b** Region enrichment results illustrating the ChromHMM annotation enrichment at loop anchors and their 2 kb interval regions. Enrichment Z-scores are plotted. Cells are separated by different shapes, and regions are distinguished by different colors. **c** Bar plot showing the fold change of CTCF binding sites enrichment at loop anchors and their 2 kb interval regions. Statistical significance was calculated with treating 2 kb interval regions as background. *P* < 0.05 is asterisked. **d** Histograms showing the distribution density of genomic distance of identified loops and chromatin interactions. **e** Fraction of stable loops and interactions. Both loop anchors and anchor pairs, interacting fragments and interaction pairs in AC and OB were compared with that in hMSC. Stable interactions and loops in hMSC were counted if they are overlapped in either AC or OB.

Most of the loops and interactions spanned a genomic distance within 1 M bp with a median up to 280 kb for loops and 200 kb for interactions (Fig. 2d and Supplementary Table 6). The similar distribution was reported in IMR90 cells^22^. Comparing to loops, a larger fraction of chromatin interactions had distance less than 50 kb, suggesting the abundant intra-gene interactions. We next compared their stability towards hMSC differentiation. We found both loops and interactions changed dramatically that only less than 20% of interactions and 30% of loops in hMSC were stable in either AC or OB (Fig. 2e). As for loop anchors and interacting fragments, the loop anchors remained changeable as less than 50% of those in AC and OB were inherited from hMSC. Contrarily, the majority of interacting fragments (~73%) in AC and OB can be found in hMSC. Additionally, about 80% of interacting fragments in hMSC were stable after differentiation (Fig. 2e), suggesting that DNA interacting property in AC and OB is maintained after cell differentiation, but the contacting fragments are selectively picked when forming loop structures in different cells, also indicating that loop structures are more changeable and able to capture more cell-specific chromatin features than interactions. Together with the fact that chromatin loops spanning longer genomic distance are capable to find more long-range regulations, we therefore focused on chromatin loops in subsequent analyses.

Thus, we found prominent enrichment of transcription regulatory signals within loop anchors, suggesting the important role of loops on gene expression regulation.

### Strengthened 3D chromatin architectures are accompanied by enhanced gene expression

We generated RNA-seq data to analyze the relationship between chromatin conformation and gene expression. After quantifying gene expression level in each cell, we found that genes residing in active A compartment had higher expression level than which in inactive B compartment as expected (Supplementary Fig. 4a). Besides, gene expression increased as the distance between promoters and interacting fragments decreased (Fig. 3a). We next identified differential expressed (DE) genes in AC and OB (Fig. 3b, left panel). About 82.73 ~ 88.14% of up-regulated genes entirely reside in TADs (Fig. 3c). We then explored the genomic position of up-regulated genes towards chromatin loops and observed strong transcription signals of up-regulated genes around the loop anchors of AC and OB (Supplementary Fig. 4b). These observations provide evidence from different Hi-C data scales that active genes usually locate at genomic area with detectable 3D structures.

**Fig. 3.**
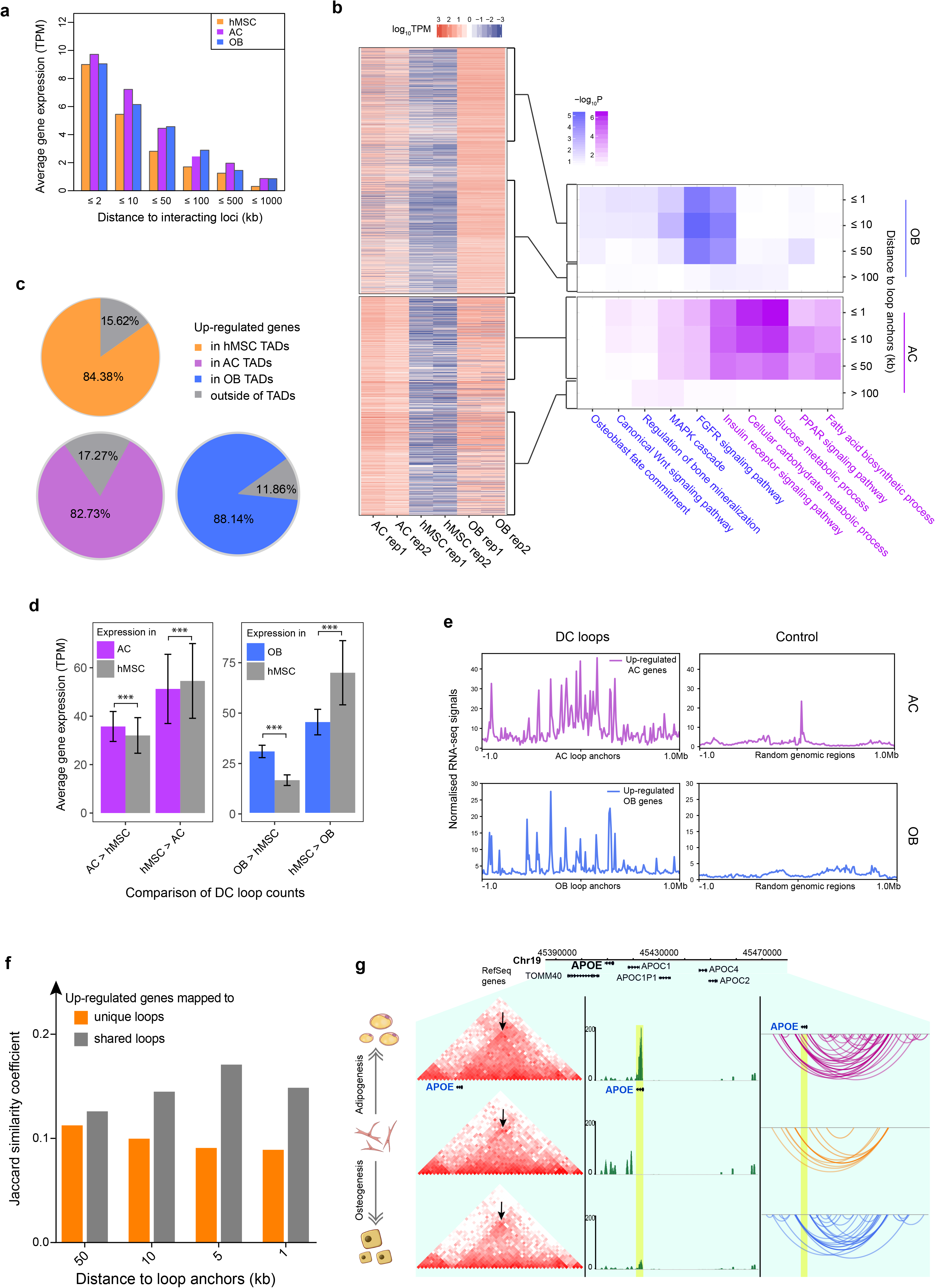
Chromatin 3D structure is coupled with active gene expression, and differentially contact loops are close related to gene activities after adipogenic and osteogenic differentiation. **a** General gene expression level with respect to different distances to interacting fragments. **b** (Left panel) Heatmap showing the Gene expression TPM of up-regulated genes in AC and OB. (Right panel) Differentiation associated GO pathway enrichment using up-regulated genes located within different distances to DC elevated loop anchors **c** Pie charts showing the fraction of up-regulated genes located in or outside of TADs. **d** Comparison of gene expression levels in AC/OB and hMSC with respect to mapping DC loop counts. *P* values were calculated by paired-sample t-test. **e** Up-regulated gene distribution along ± 1 Mb flanking regions of AC/OB loop anchors and randomly selected genomic regions that match the length and GC content of loop anchors. RNA-seq signals were RPKM normalized. **f** The Jaccard similarity coefficients indicate the gene set similarity between AC and OB with genes mapping to either unique or shared loops under different genomic distance to loop anchors. **g** An illustration of differential contact loops near *APOE* with enhanced contact frequency (heatmap, left) and increased pairwise interactions (arc, right) in AC cells comparing with hMSC and OB, which is accompanied by elevated *APOE* expression level (middle). The arrow heads indicate the loop interacting position.

We next sought to investigate whether chromatin structure alternation was associated with differential gene expression. Differentially contacted loops (DC loops) were found through a statistical identification approach (see Methods). After correcting with 5% FDR, 6,889 and 7,031 elevated DC loops were identified in AC and OB, respectively (hereafter refer to as AC/OB loops). We then counted nearby DC loops (± 1 Mb around TSS) for each gene, and separated genes by loop counts difference between cells. As shown in Fig. 3d, gene expressions were significantly higher in cells having more DC loops (Wilcoxon signed-rank test). Furthermore, as up-regulated genes usually contain key regulators for cell differentiation, we explored the genetic position of these genes towards DC loops. The RNA-seq data of up-regulated genes showed strong transcription signals around AC and OB loop anchors, while as a comparison, were less obvious at random length and GC content matching regions (Fig. 3e). These suggest the colocalization of active genes and strengthened contacting regions.

Upon further diving into DE genes in AC and OB, we measured the distance between up-regulated gene TSSs and the nearest DC loop anchors. Genes under different distance to anchors were gathered to perform pathway enrichment analysis. We found that as the distance decreased, the enrichment of adipogenesis and osteogenesis related pathways increased. Especially, when under 50 kb, a significant proportion of up-regulated genes in AC and OB started to be involved in the biological process of cell function determination (Fig. 3b, right panel). Notably, significant enrichment under 1 kb distance indicated the connections of gene bodies/promoters with distal fragments, suggesting the possible cases of long-range promoter-enhancer regulation. Genes as cell fate determinants are supposed to be cell-specific activated, which, to our anticipation, should be partially modulated by cell-specific loop formation. Therefore, we found exclusive “unique loops” by comparing AC and OB loops, and loops with both anchors overlapping were referred to as “shared loops”. 4,899 and 5,062 unique loops were found in AC and OB, respectively. We next mapped up-regulated genes to either loops at different distance cut-off.

Genes were then tested for similarity between AC and OB (Jaccard similarity coefficient; Fig. 3f). The results showed that gene similarity under 50 kb distance were comparable between unique and shared loops. However, when the distance decreased, gene similarity decreased at unique loops, while increased at shared loops. This suggests the association between cell-specific gene activation and exclusive loop formation that is close to gene. Take obesity gene *APOE* as an example, which encodes a major protein of the lipid and lipoprotein transport system. The −1.01 kb upstream region from *APOE* was interacting with a downstream fragment forming a 50 kb loop. This unique AC loop was confirmed by the increased contact frequency and chromatin interaction in AC (Fig. 3f). As expected, the gene expression profile showed that *APOE* was specially activated in AC after differentiation. This evidence links cell-specific gene activation with unique loop formation.

### Strengthened loops after differentiation are associated with enhancer generation

Distal enhancer-promoter contact is one of the most important features for loop formation^29, 30^. We next intended to investigate whether differential gene expression was caused by rearrangement of enhancer-promoter interactions. We initially generated ChIP-seq data of two enhancer-associated markers in 3 cells, histone H3 lysine 4 monomethylation (H3K4me1) and lysine 27 acetylation (H3K27ac). By computing and integrating peaks, we identified 128,179, 224,322 and 167,451 putative enhancers in hMSC, AC and OB, respectively (see Methods). We found limited number of shared enhancers between hMSC and AC (9.84%) or OB (12.15%), and the rest were considered as “differential enhancers”. The correlation test suggests that both histone signals are correlated between replicates but are cell-specific across cells (Fig. 4a).

**Fig. 4.**
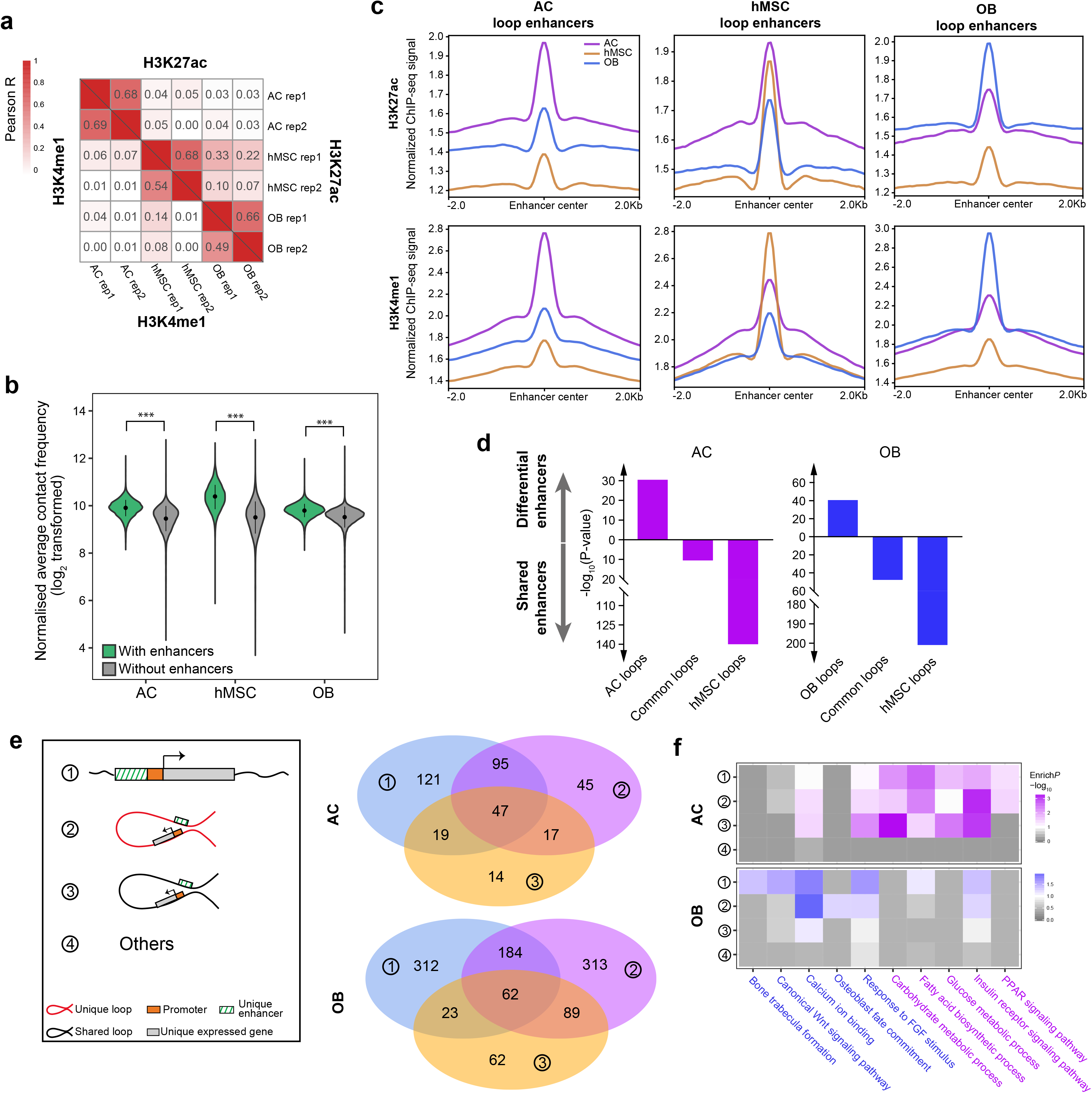
Adipogenesis and osteogenesis are achieved by strengthened loops featured with cell-specific enhancers. **a** Heatmap of Pearson correlation coefficients for H3K27ac and H3K4me1 ChIP-seq signals. **b** Violin plot showing the contact frequency difference between loops with and without enhancer mapping. Statistical significance was calculated by t-test. **c** Comparison of H3K27ac and H3K4me1 distribution at cell-specific loop enhancers (CSLE) among 3 cells. ChIP-seq signals are normalized for reads count and length. **d** Enhancer enrichment at DC loops. Fisher exact test was used to determine enrichment status, and enrichment direction was defined by odds ratio. **e** Diagram showing 4 proposed transcriptional regulation patterns (left panel), and the pathway enrichment with unique expressed genes in each pattern (right panel). GO pathways related to adipogenesis or osteogenesis are coded by different colors. **f** Venn diagrams showing the gene overlapping between different patterns.

We then mapped putative enhancers to chromatin loops. The contact frequency of loops mapping with enhancers was significantly higher than loops without enhancers mapping (Fig. 4b), which was in accordant with the mechanism that enhancer-mediated interaction is associated with strengthened chromatin contact. To test whether DC loops were prone to harbor differential enhancers, we picked out enhancers that located in DC loops as “loop enhancers”. The H3K4me1 and H3K27ac signals on these enhancers from different cells showed that both histone marks from AC and OB had exceeding signals in the same cell as loop enhancer annotated, while histone marks from hMSC loop enhancers were appeared to be shared across cells (Fig. 4c). This result hints that some enhancers in AC and OB could be inherited from hMSC, but enhancers in DC loops after differentiation are more cell specific. In addition, we also performed statistical enrichment analysis to see where the differential enhancers prefer to locate (see Methods). Consistently, AC and OB loops were significantly enriched with differential enhancers (fisher exact *P* = 3.22× 10^−31^, OR = 1.20 for AC; *P* = 2.61× 10^−41^, OR = 1.25 for OB), while both common and hMSC loops were enriched with shared enhancers (Fig. 4d). Thus, the significant enrichment of differential enhancers in AC and OB loops links novel enhancer generation with loop formation after differentiation.

### Cell fate determination is achieved by unique loops mapping with cell-specific enhancers

We next were interested to know what role the enhancer may play in cell fate determination. We focused on unique enhancers and unique genes in AC and OB, which were exclusive enhancers and up-regulated genes comparing the other two cells. We hypothesized 4 enhancer-mediated regulation patterns resulting in elevated gene expression after differentiation (Fig. 4e, left panel). The first was regulated by unique enhancers located at unique gene promoters (upstream 5 kb from TSS). The second and third were both related to distal enhancer-promoter interactions but in unique loops and shared loops, respectively. The other situations were considered as the fourth pattern. Unique genes were mapped to each pattern according to their genetic locations. By counting unique gene numbers in each pattern, we observed that up to 57% of direct enhancer mapping genes were also undergone putative long-range regulation (Fig. 4e, right panel). Considering the majority of genes overlapping between 2D and 3D regulation, we then would like to know which one should play a predominant role. Previous study has shown that genes within a TAD are more co-expressed than do those in different TADs^31^. We therefore hypothesized that the intra-loop enhancers are able to synchronously regulate the genes locating in the same loop through chromatin looping, and by this way, leads to co-expression, even though the enhancers are mapped to their local genes (referred to as “tag genes”). Therefore, we explored GTEx data (phs000424.v8.p2) to perform expression correlation between tag genes and other genes within the same loops. We used adipose as the target tissue for AC. Because the link between blood and bone biology has been widely discussed^32, 33^, we used whole blood as the target tissue for OB. We generated the background set in each tissue comprised of pairwise genes that were randomly selected from genome-wide, and performed correlation tests as well. The same length of multiple tests were conducted. The comparison of correlation coefficient (Pearson’s R) showed that genes located within the same loop were more co-expressed than background (t-test, *P* < 0.05; Supplementary Fig. 5), supporting the enhancer synchronous regulation by taking advantage of chromatin loops.

Pathway enrichment analysis was then performed with unique genes in each pattern. The results recognized significant enrichment of adipogenesis and osteogenesis related pathways with genes directly mapped with unique enhancers or manipulated by putative long-range regulation in unique loops (Fig. 4f). The canonical enhancer adjacent to promoter directly regulates gene expression, which is the most acceptable way to manage cell determination. Here we showed that the 3D chromatin regulation function was also noticeable in favor of adipogenesis and osteogenesis. Intriguingly, adipogenesis pathways specially contained genes located within shared loops, however, we didn’t see such preference of osteogenesis pathways (Fig. 4f). This hints that some genes controlling adipogenesis lies in shared loops, and are adipocyte-specifically activated by the manipulation of particular enhancers. We observed the importance of insulin receptor signaling pathway for OB, which controls osteoblast development through stimulating osteocalcin production and suppressing Twist2^34^, and the enriched process of “response to FGF stimulus” in AC, which can be explained by favoring adipose tissue development and metabolism by several FGF family members^35^.

Taken together, we highlighted the important role of enhancer-centered regulation within chromatin loops, and cell-specific loops accompanied with cell-specific enhancers after hMSC differentiation are crucial to cell fate commitment.

### Chromatin accessibility reveals loop-associated cell-specific regulator activation

Chromatin accessibility is a critical condition for enhancer-anchored gene regulation. Thus, we identified chromatin accessible regions in AC and OB using ATAC-seq. Correlation analysis validated replicates concordance (Supplementary Fig. 6a). We finally identified 138,820 and 120,209 confidential peaks in AC and OB, respectively. We tested the colocalization and correlation between enhancer and chromatin accessibility, and found that H3K27ac was highly correlated with ATAC-seq peaks (Supplementary Fig. 6b). Comparing with randomly selected regions (with GC and length matched), ATAC-seq peaks were successfully colocalized with H3K27ac marking regions (Fig. 5a). Notably, the ATAC-seq peak summits occur at H3K27ac depletion sites, which is identified as the available regions for TFs binding^36^. As the open accessibility at promoters, we detected accessible chromatin around promoters of up-regulated genes, and found that 84.71% and 68.82% of those in AC and OB were mapped with reliable ATAC-seq signals (peak filtering *P* < 0.05). We next mapped ATAC-seq peaks to gene promoters and estimated the contact frequency within each peak as well as the corresponding gene expression levels. Using matched random regions as control, we confirmed that open chromatin is significantly associated with higher contact density and gene expression levels (Fig. 5b). This suggests the easier accessibility at chromatin interacting regions, which is essential to make DNA available for regulatory factor binding in favor of gene activation.

**Fig. 5.**
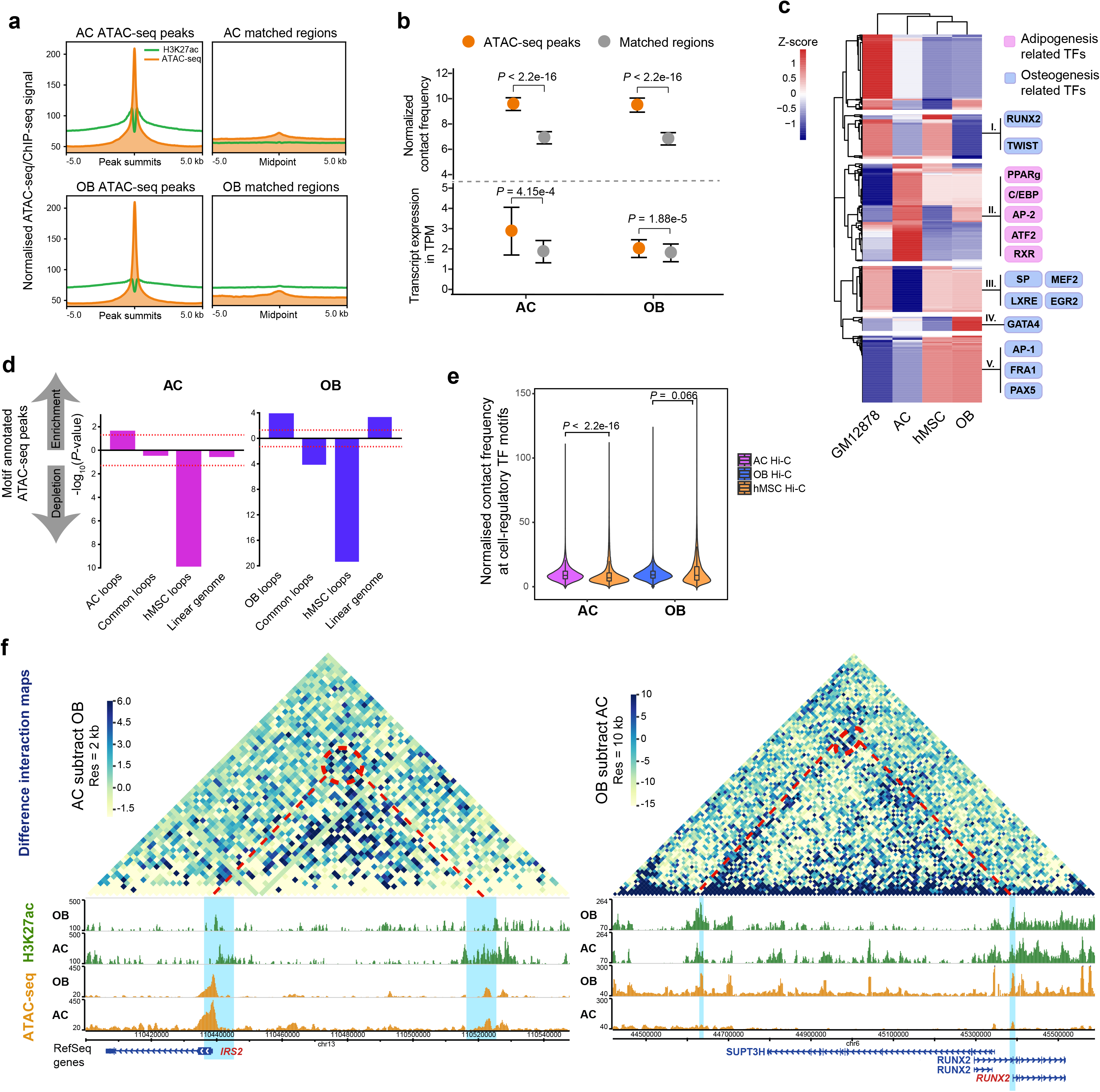
Chromatin accessibility reveals loop-mediated transcription network reprogramming after hMSC differentiation. **a** Colocalization of chromatin accessible and H3K27ac modification regions. The colocalization was compared between ATAC-seq peaks (left column) and randomly selected regions with length and GC content matched regions (right column). **b** Comparison of normalized contact frequency (top) and expression level of promoter mapping transcripts (bottom) between ATAC-seq peaks and randomly matched regions. Statistical significance was estimated by t-test. **c** Heatmap showing the enrichment Z-scores of 413 known motifs at chromatin accessible regions in each cell type. Known regulatory TFs are listed on the right with separating adipogenesis and osteogenesis related TFs by different colors. **d** Chromatin structure enrichment of ATAC-seq peaks mapping with cell-regulatory motifs. Fisher exact test was used to determine enrichment status, enrichment direction was defined by odds ratio. Red dashed line indicates significant threshold *P* = 0.05. **e** Violin plot indicating the difference of normalized Hi-C contact frequency at cell-regulatory motifs between hMSC and differentiated cells. Statistical significance was estimated by t-test. **f** Heatmaps of subtractive interaction matrix and genome browser screenshots showing the unique loop structure and differential enhancer and open chromatin signals for *IRS2* and *RUNX2*.

We further focused on interrogating whether loop formation is coupled with activating particular TF binding affinity after hMSC differentiation. We retrieved ATAC-seq data of hMSC and GM12878 from ENCODE^37^, and identified motifs enriched in open chromatin regions in each cell using the other 3 cells as background (see Methods). All known TF motifs were clustered and stratified with respect to their enrichment Z-scores (Fig. 5c). Notably, we predicted distinguished TF bindings in AC, however, some putative TF motifs had comparable Z-scores between OB and hMSC. We next classified 5 TF groups according to their cell-specific activation manner. Known adipogenesis associated TFs, like PPARg, C/EBP and AP-2, were successfully found in AC-specific motif group. Interestingly, some osteogenesis related TF motifs were accessible in both hMSC and OB cells (Fig. 5c). The TFs distinction between adipogenesis and osteogenesis has been reported by Rauch *et al.* In their paper, they found *de novo* TF activation during adipogenesis, while activation of MSC TFs is required in response to osteogenic stimulation^15^, which is in agreement with the difference of TF clusters showing here. These cellular active motifs were collected as “cell-regulatory TF motifs”. Next, to find out which genomic position is prone to contain cell-regulatory TF motifs, we performed enrichment analysis in DC loops and linear chromatin synchronously (see Methods). Chromatin accessible regions with AC-regulatory and OB-regulatory TF binding were significantly enriched in AC loops and OB loops, respectively (Fisher test, *P* < 0.05; Fig. 5d). On the other hand, the results indicated that the strengthened loops in hMSC were less involved in cell differentiation. Considering the differentiation related genes usually remain inactive/poised in hMSC, this observation points out a link between gene inactivity and loop disconnection. The linear chromatin regions showed particular enrichment with OB-regulatory TF motifs (Fisher test, *P* < 0.01; Fig. 5d), suggesting that, to some extent, osteogenic differentiation are also modulated by 2D regulation. This might be the reason that hMSC and OB share some effective TFs. In addition, we retrieved Hi-C data at these cell-regulatory TF motifs to explore their contact frequency. We detected more frequent chromatin contact at the AC-regulatory TF motifs in AC cell than that in hMSC cell, which significantly passed the statistic test (U test, *P* < 2.2×10^−16^; Fig. 5e). As for OB-regulatory TF motifs, however, the comparison showed a trend for significant difference between OB and hMSC (U test, *P* = 0.066; Fig. 5e), which can be explained by the enrichment results that part of these putative TF binding sites particularly locate at genomic regions free from spatial interaction in both cells.

Next, to investigate how the accessible regulatory TF motifs are involved in long-range regulation of adipocyte and osteoblast commitment, we selected unique AC/OB loops with unique enhancer and gene promoter interaction, followed by mapping enhancers and promoters to AC and OB regulatory motifs separately. By this way, we screened out 25 and 40 unique expressed genes in AC and OB, respectively (Supplementary Table 7). Among them, we successfully recognized adipocyte functional genes *PDK4*^38^, *IRS2*^39^, *SULF2*^40^, *PTGS1*^41^ etc. and osteoblast functional genes *RUNX2*^42^, *SIGLEC15*^43^, *SCMH1*^44^ etc. The genome browser illustrations for *IRS2* and *RUNX2* (Fig. 5f) show unique loops connecting distal unique enhancers with promoters of candidate transcript, and meanwhile, the differential ATAC-seq signals make both enhancer and promoter accessible to achieve cell-specific gene activation, and eventually promote cell commitment.

Together, we revealed a close connection between accessible chromatin and loop formation after hMSC differentiation. Moreover, coupling with cell-specific enhancer mapping, this connection is important for cell-determined gene activation under long-range chromatin regulation.

### Comprehensive loop-mediated regulatory networks indicate key regulators for adipogenesis and osteogenesis

So far, we have emphasized the association between chromatin loops with gene regulation by mapping different regulatory elements. We next sought to construct regulatory networks to tie multi-omics data together and find out the prospective loop-mediated regulation cascades for adipogenesis and osteogenesis. The network was constructed based on unique AC/OB loops. We assumed that these loops shorten the spatial distance between distal enhancers and target genes in a cell-specific manner, which as a result, makes TFs easily bind to target genes. Fig. 6a shows the regulatory model that is also our strategy to build up networks. Briefly, the ATAC-seq peaks containing cell-specific regulatory TF motifs were first used to screen loops with both accessible anchors. Loop anchors were next mapped with promoters of gene transcripts at the one side and the unique enhancers at another side. TF binding events retrieved from GTRD^45^ were used to identify TFs binding at both enhancers and promoters (see Methods). After weighting TFs and target genes with expression fold change, focusing on unique expressed genes, and filtering by weight > 1, we identified 23 and 38 genes involved in loop-mediated regulatory networks in AC and OB, respectively (Fig. 6b; Supplementary Table 8). Among 14 and 19 genes in AC and OB regulatory networks that can be found detectable mutated mouse phenotypes from Mouse Genome Informatics (MGI) database^27^, 8 and 6 genes were annotated with adipose and skeleton relevant disfunctions, respectively (Supplementary Table 9). Among all gene nodes, *CELSR1* and *PRLR* are linked to 55 and 52 TFs, which are the maximum number of TF annotations among genes in AC and OB regulatory networks. Mouse with deficient *CELSR1* and *PRLR* gene are associated with decreased body size and decreased bone mass, respectively (Supplementary Table 9), emphasizing the important biological functions of these two genes. Among all TF nodes, the ones that involved in adipogenesis and osteogenesis had more abundant edges than others (Fig. 6b). Two representative genes are *IRS2* and *RUNX2* that are essential to adipocyte^39^ and osteoblast^42^ differentiation, respectively. We successfully detected an unique AC loop anchored at *IRS2* promoter and interacted with a distal enhancer (with 90 kb chromatin interval) (Fig. 6d). We showed the possible binding of 20 TFs at both *IRS2* promoter and distal enhancer (Fig. 6c, upper panel), and intriguingly, we found the particular binding affinity of two genome architectural proteins CTCF and YY1, supporting the long-range interaction. We have revealed the dominant ATAC-seq and H3K27ac signals focusing *IRS2* (Fig. 5f), here, we confirmed the CTCF binding at both anchors containing *IRS2* promoter and distal unique enhancer by obtaining ChIP-seq data from human adipose (GSE105994)^37^ (Fig. 6d, upper panel). The prominent *IRS2* expression in AC reflects the cell-specific activation consequence. Similarly, we found a regulatory network especially connecting a *RUNX2* transcript promoter with a 770 kb interval region whose cell-specific accessibility and enhancer property were indicated through ATAC-seq and enhancer mapping (Fig. 5f). The TFs screening further suggested 9 binding events (Fig. 6c, lower panel). Previous studies have characterized the irreplaceable role of ESR1 on osteoblast development^46^. We therefore obtained ESR1 ChIP-seq data from osteosarcoma cell (GSE26110)^47^. The ESR1 peaks were appeared not directly at but closely to both loop anchors (Fig. 6d, lower panel), which should be explained by the frequent occupation of structural proteins at loop anchors. Here, we demonstrated that the regulatory network is robust to illustrate loop-mediated gene regulation. We believe that our strategy is also applicable for finding novel target genes functioning as cell fate determinant during hMSC differentiation.

**Fig. 6.**
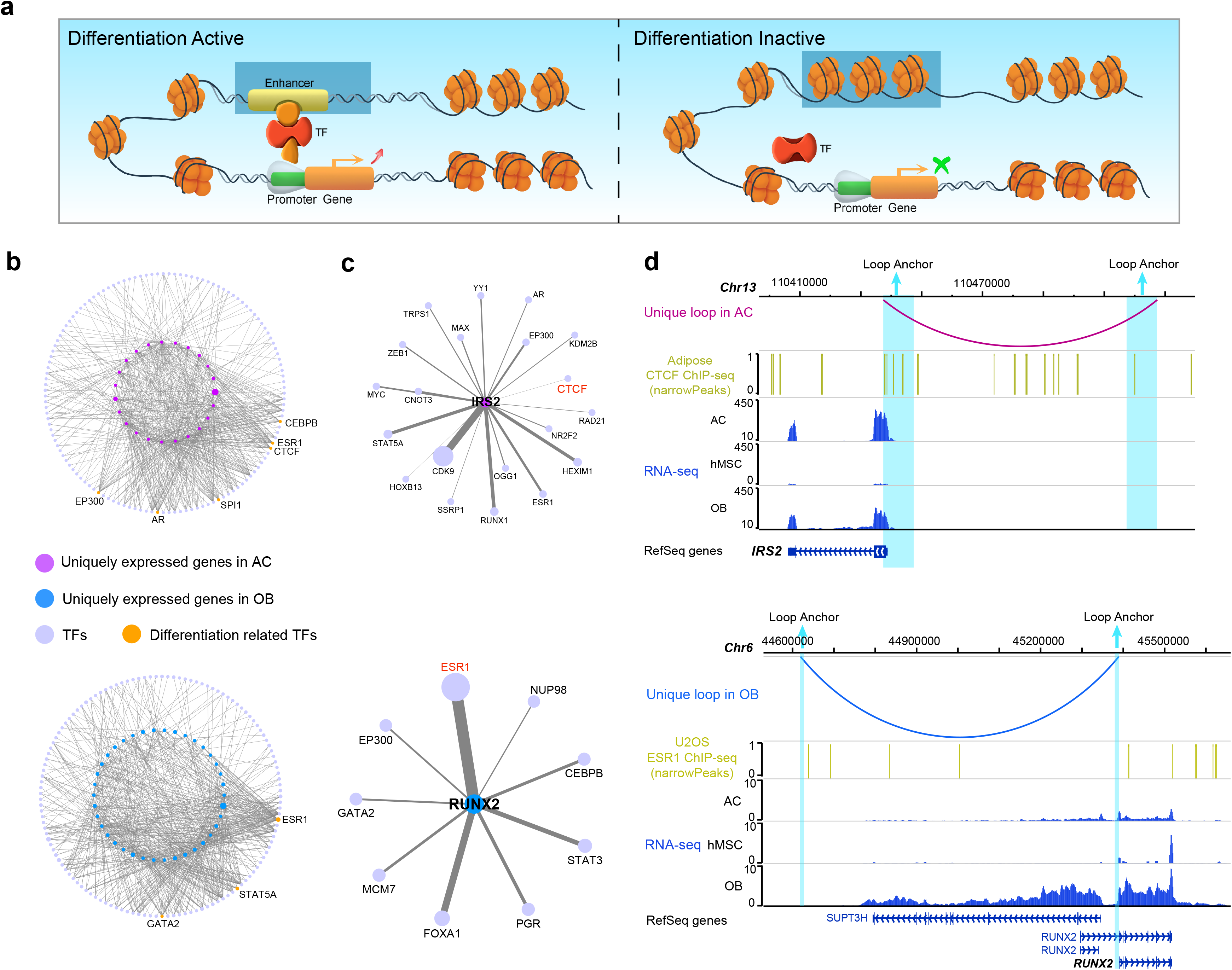
Regulatory networks identify loop-mediated gene regulation cascades for cell fate determination. **a** The network construction strategy illustrating the mechanism of gene activation to achieve cell-type commitment through the spatial proximity of long-range promoter and unique enhancer facilitating by unique loops. **b** Regulatory networks targeting unique expressed genes in AC and OB. Adipogenesis and osteogenesis related TFs shown names aside are marked in orange. **c** Representative networks for adipogenesis related gene *IRS2* and osteogenesis related gene *RUNX2*. **d** Genome browser screenshots showing the unique loop structure and cell-specific gene expression for adipogenesis related gene *IRS2* and osteogenesis related gene *RUNX2*. The ChIP-seq data of putative TF CTCF ESR1 were added to show the expected binding sites around loop anchors.

### eQTL variants are linked with target genes through chromatin loops

We next were interested in whether eQTLs can be linked with target genes through loop formation. We included eQTL data from adipose and blood-derived tissues from GTEx and mapped snp-gene pairs to AC or OB loops. The Q-Q plots indicated a superior significance of eQTL associations at AC/OB loops than that at hMSC loops (Fig. 7a; Supplementary Fig. 7), and the dominance was even more obvious than eQTLs without loop mapping (Kolmogorov-Smirnov test *P* < 2.2 × 10^−16^). This indicates that the eQTLs supporting by DC loops have stronger association with gene expression, and confirms the loop-mediated regulation mechanism. Hence, in light of the eQTL dominance at DC loops, we added eQTL information to regulatory networks. The SNPs locate at TF binding sites and impact target gene expression were linked to the networks. Eventually, we suggest 14 and 20 genes that are possibly activated by loop-mediated regulatory cascades in AC and OB, respectively (Supplementary Table 10). Particularly, we identified 5 and 2 SNPs (pruning with LD < 0.8), recognized as the eQTL sites for *IRS2* and *RUNX2*, potentially interrupt TFs binding and impact the long-range regulation (Fig. 7b).

**Fig. 7.**
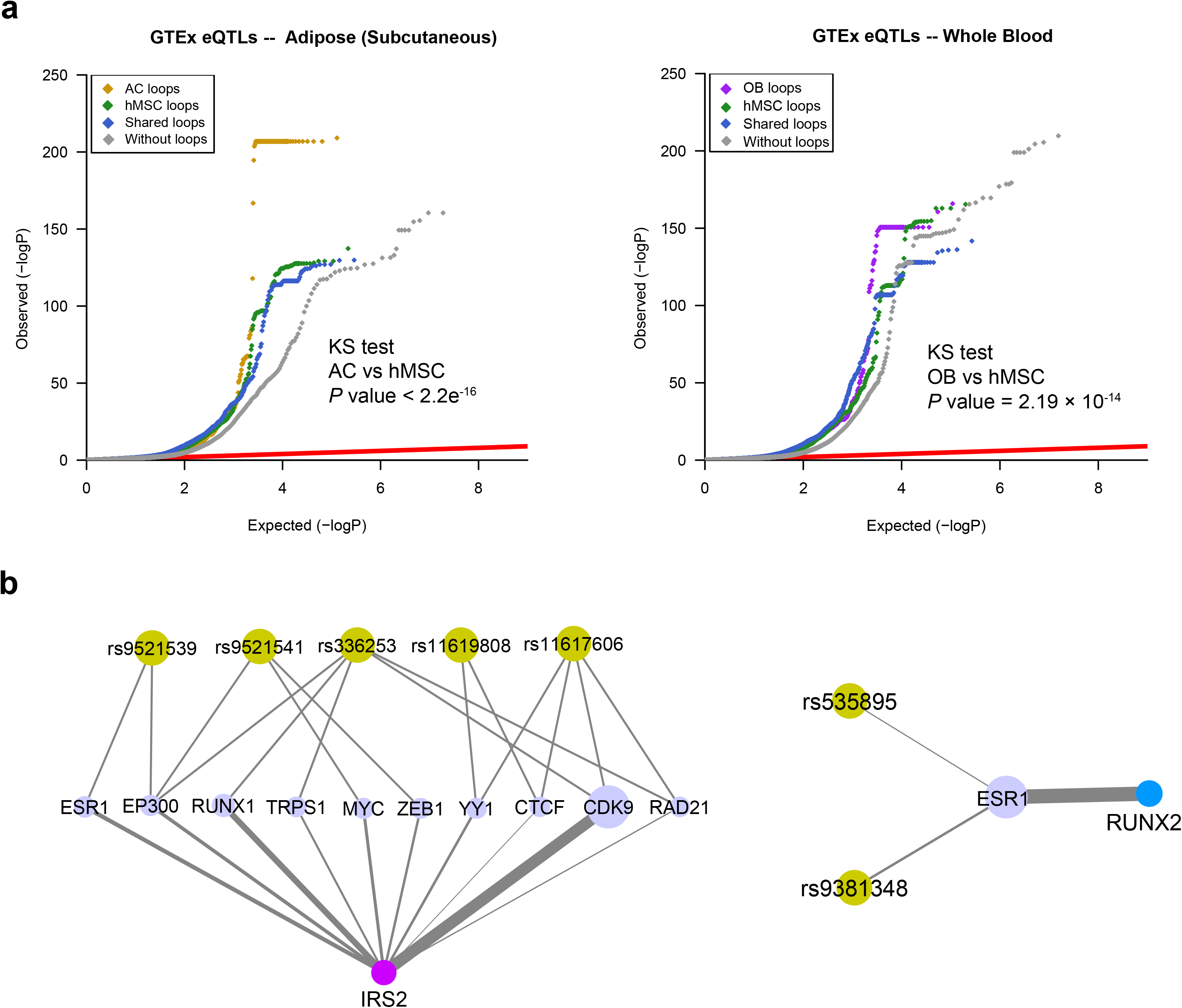
eQTL variants are linked to target genes through chromatin loop structures. **a** Q-Q plots drawn with eQTL data from subcutaneous adipose (left panel) and whole blood (right panel) tissues illustrating the significant enrichment of eQTL associations at AC/OB loops. **b** The regulation cascades identified for *IRS2* and *RUNX2* through jointly analyzing multi-omics data. eQTL snps were marked in yellow-green, and the putative TFs were marked in lavender.

Taken together, we have revealed the associated loop changes after inducing hMSC to adipogenesis and osteogenesis differentiation. We also suggested a close relationship between cell-specific loops and adipocyte/osteoblast determination, which is expected to provide better understanding of the controlling factors for hMSC differentiation.

## Discussion

Here, taking advantage of high resolution Hi-C data, we’ve recognized chromatin loops for hMSCs and the differentiated adipocytes and osteoblasts. We also performed the comprehensive assessment of mRNA expression, histone modification as well as chromatin accessibility. After leveraging these data, we identified differential contact loops in each cell and screened out unique loops for differentiated cells. Subsequentially integrative analyses linked strengthened loop formation to gene activation and suggested significant enrichment of differential enhancers and TF motifs at strengthened loops. Furthermore, we linked unique loops with cell-specific enhancers and accessible TF motifs, and identified the robust long-range gene regulation mechanism responsible for cell fate determination. Finally, we constructed regulatory networks involved in adipocyte and osteoblast commitment and emphasized the loop-mediated regulation cascades especially for *IRS2* and *RUNX2* that leads to adipogenesis and osteogenesis, respectively. Overall, our study provides the first investigation of 3D chromatin structure changes after hMSC were stimulated to adipogenic and osteogenic differentiation. According to the robust analytical evidence, we emphasize the long-range regulatory mechanisms for hMSC differentiation.

The cell fate determination during hMSC differentiation requires cell-specific genes activation or repression. Here, we focused on active regulatory elements attempting to reveal the mechanisms that underlie the gene activation after hMSC responding to differentiation induction. Although we rarely mentioned whether differentiation-repressed genes are inhibited by prohibiting loop interaction during this process, the evidence that highly expressed genes in hMSC had lower interaction strength in differentiated cells shows the inhibition of genes by disrupting the 3D structures during cellular function alternation. Further investigation towards these genes is needed to confirm their effect on hMSC function maintenance.

## Methods

### Cell culture and hMSC differentiation

Primary human umbilical cord derived hMSCs were obtained frozen from Shaanxi Stem Cell Engineering Co., Ltd from 1 donor who have signed the informed consent for this study. Cells were thawed and expanded for an additional passage for all the subsequent experiments. hMSC cells were seeded at a density of 1 × 10^4^ cells/cm^2^ and cultured at 37 °C, 5% CO_2_ in Dulbecco modified Eagle medium (DMEM; GE) supplemented with 10% fetal bovine serum (FBS; GIBCO) and 1% antibiotics (penicillin 100 U/ml, streptomycin 100 μg/ml; Solarbio Co., Ltd). When 80% confluence was reached, part of the cells was harvested, and the left were switched to differentiation culture medium to induce adipogenesis and osteogenesis.

For osteoblastic differentiation, hMSC cells were grown in DMEM medium supplemented with 10% FBS, 1% penicillin/streptomycin, 10 mM glycerol-2-phosphate (Sigma), 50 μM L-ascorbic acid (Sigma), and 100 nM dexamethasone (Sigma) for 21 days. Medium was replaced every 3 days.

Adipogenic differentiation was induced in hMSC cells cultured by alternately supplying treatment of solution A and B. Solution A: DMEM medium containing 10% FBS, 1% penicillin/streptomycin, 10 mg/L insulin (Novo Nordisk), 1 umol/L dexamethasone, 0.5 mmol/L IBMX (Sigma), 100 umol/L indometacin (Sigma). Solution B: DMEM medium containing 10% FBS, 1% penicillin/streptomycin, 10mg/L insulin. Cells were firstly cultured in solution A for 3 days and were additionally supplied with solution B for another day. Cells were harvested after adipogenic induction for 15 days.

Cell differentiation status were further verified at 4 time points (0d, 5d, 10d, 15d for adipogenic differentiation; 0d, 7d, 14d, 21d for osteogenic differentiation) through microscopic identification, Oil Red O/Alizarin Red S staining and qRT-PCR quantification of marker genes (Supplementary Fig. 1). The staining areas were counted by ImageJ^48^ software at each time point, and the statistical significance was indicated by t-test.

### Hi-C library preparation and sequencing

6 technical replicates of adipocytes and osteoblasts, and 7 technical replicates of hMSC were generated after cell differentiation with each replicate containing about 1 × 10^7^ cells. In situ Hi-C was next performed on each replicate using methods as previously described^26^. Briefly, after harvesting from plates, cells were crosslinked with 1 ml of freshly made 1% formaldehyde solution and incubated for 10 min at room temperature. The reaction was quenched by adding glycine solution to a final concentration of 0.2 M. Cells were lysed and chromatin was next digested with 200 U of MboI restriction enzyme for 16 h at 37 ◻. Digested DNA ends were labeled using biotinylated nucleotides and incubated at 37 ◻ for 90 min. Fragments were proximity ligated by adding T4 DNA ligase and were incubated at 4 ◻ for 1 h, followed by 4 h at room temperature. Samples were supplemented with SDS, Proteinase K, and NaCl to reverse crosslinking, and incubated overnight at 65 ◻. After that, DNA fragments were purified and dissolved.

Purified DNA fragments were sheared to a size of 300-500 bp. Ligation junctions labeled with biotin were subsequently pulled down using streptavidin C1 beads and prepared for Illumina sequencing. TA cloning was carried out to examine the library quality. Hi-C libraries were sequenced on an Illumina HiSeq X Ten system. The Hi-C experiment and library sequencing were performed by Novogene Co., Ltd, Beijing, China.

### RNA-seq data generation

Two technical replicates were generated for each cell type. Total RNA was extracted form samples using the TRIzol (Invitrogen) method^49^. RNA concentration and purity were evaluated with a NanoDrop spectrophotometer (Thermo Fisher). 6 libraries were constructed under manufacturer’s instructions and were then sequenced on the Illumina HiSeq X Ten platform using the 150-bp pair-end sequencing strategy. Finally, an average of 47 M pair-end reads were obtained per sample.

### Chromatin immunoprecipitation assay

ChIP assay was performed using the SimpleChIP Enzymatic Chromatin IP Kit (Cell Signaling Technology) as previously described^50^. Briefly, cells were crosslinked with 1% formaldehyde. After quenching with glycine solution, cells were rinsed, pelleted and resuspended in cold PBS, and then resuspended and pelleted twice with buffer A and B, respectively. Nucleus were digested with Micrococcal Nuclease (2,000 gel units/μL). The digestion reaction was deactivated with 0.5 M EDTA. The nucleus were then pelleted, and sediment resuspended in ChIP buffer using protease inhibitor cocktail. The lysate was sonicated with the VirTis Virsonic 100 Ultrasonic Homogenizer/Sonicator for 3 pulses. After centrifuging, the supernatant was collected and was immunoprecipitated with H3K4me1 and H3K27ac antibodies (Abcam) or normal immunoglobulin G (IgG) as a negative control, and precleared with agarose beads. DNA protein complex was then precipitated with agarose beads, eluted from the beads, and reversely cross-linked by 5M NaCl and Proteinase K. Libraries for ChIP-seq were prepared following Illumina protocols. Libraries were next sequenced on the Illumina HiSeq X Ten platform configured for 150-bp pair-end reads.

### ATAC-seq data generation

ATAC-seq libraries were constructed for adipocytes and osteoblasts following the original protocol^51^. In brief, two hundred thousand cells were lysed with cold lysis buffer (10◻mM Tris-HCl, pH◻7.4, 10◻mM NaCl, 3◻mM MgCl2 and 0.03% Tween20), and centrifuged at 500g for 8◻min at 4◻°C. The supernatant was carefully removed, and the nuclei was resuspended with Tn5 transposase reaction mix (25◻μl 2◻×◻TD buffer, 2.5◻μl Tn5 transposase and 22.5◻μl nuclease-free water) (Illumina) at 37◻°C for 30 min. Immediately after the transposition reaction, DNA was purified using a Qiagen MinElute kit. Libraries were sequenced on an Illumina HiSeq X Ten sequencer to an average read depth of 52 million pair-end reads per sample. The ATAC-seq experiment and library sequencing were performed by Frasergen Bioinformatics Co., Ltd, Wuhan, China.

### qRT-PCR

Cells at each differentiation time point were partially collected to detect marker gene expression. Total RNA was isolated with Trizol reagent (Invitrogen), and was converted to cDNA with reagents purchased from Vazyme Biotech Co., Ltd. PCR procedure was performed using Qigen SYBR Green PCR Kit (Qiagen) and was operated with Bio-Rad System (CFX Connect™, Bio-Rad). The following specific oligonucleotide primers were used: *PPARG* (5’- AGCCTCATGAAGAGCCTTCCA, 3’- TCCGGAAGAAACCCTTGCA), *CEBPD* (5’- GGTGCCCGCTGCAGTTTC, 3’- CACGTTTAGCTTCTCTCGCAGTTT), *ALPL* (5’ - CCTGCCTTACTAACTCCTTAGTGC, 3’ - CGTTGGTGTTGAGCTTCTGA), *RUNX2* (5’ - GCGCATTCCTCATCCCAGTA, 3’ - GGCTCAGGTAGGAGGGGTAA), *BGLAP* (5’ - AGCGAGGTAGTGAAGAGAC, 3’ - GAAAGCCGATGTGGTCAG), *COL1A1* (5’ - TTTGGATGGTGCCAAGGGAG, 3’ - AGTAGCACCATCATTTCCACGA). All the experiments were conducted following the manufacturer’s instructions.

## Computational analysis

### Hi-C data processing

Hi-C reads from each replicate were aligned (hg19), filtered and paired using HiCUP pipeline^24^ with parameters (–longest 800 –shortest 150). In summary, ~0.53 B (~81% of total read pairs) paired reads uniquely mapped to the genome. After removing self-ligation and invalid pairs, ~0.46 B (~61% of total read pairs) valid pairs were remained for subsequent analysis (Supplementary Table 1). Valid pairs for replicates of each cell type were combined to generate raw contact matrices at different binning resolutions. We next normalized the raw contact matrices using ICE normalization^52^ with parameters (--filter_low_counts_perc 0.02 --eps 0.1 --remove-all-zeros-loci). To evaluate Hi-C data reproducibility, interacting counts at each bin were retrieved at 40 kb resolution, and Pearson correlation test was implemented between replicates.

### TAD calling and TAD signal calculation

TADs were called with Domaincalling pipeline as Dixon et al. described^17^. The ICE-normalized matrix was subjected to calculate DI (Directionality Index) values, and the results as input were applied with Hidden Markov Model (HMM) model to call TADs. We executed this TAD calling procedure at 40 kb binning resolution in this study. In total, 3,142, 2,854 and 4,968 TADs were identified for hMSC, adipocytes and osteoblasts, respectively.

To compare TAD structure between different cells, we evaluate TAD signals that indicate the strength of TAD contact. We used the method described by Ke et al. ^16^ to calculate TAD signals. First, Intra-chromosomal maps were prepared at 40 kb resolution. The TAD signal for each bin was next calculated as the log2 ratio of the number of normalized upstream-to-downstream interactions within a 2 Mb region. Bins with less than 10 counts within either upstream or downstream region were filtered. TAD signals were then used to perform Pearson correlation test between cell types.

### Identification of A/B compartments and chromatin interactions

Hi-C output from HiCUP was transformed to compatible file format to work with HOMER software^53^. We next used HOMER to calculate PC1 values and identify significant chromatin interactions. A/B compartments were determined using the “runHiCpca.pl” function with the parameters (-res 25000 -window 25000 -pc 1). The signs of the PC1 values were used to assign the chromatin into A compartment (positive PC1 values) and B compartment (negative PC1 values). For chromatin interactions, we used “findHiCInteractionsByChr.pl” function to search for pairs of fragments that have a greater number of Hi-C reads than expected by chance. Interactions were searched with the parameters (-res 1000 -superRes 2000 -maxDist 2000000). The significant interactions were identified by FDR q < 0.05.

### Identification of chromatin loops

Loops were called by two computational strategies. The first is “findTADsAndLoops.pl” function packaged in HOMER. It was utilized to call loops at 2kb resolution with parameters (-res 2000 -window 2000 -minDist 6000 -maxDist 1000000). The other software, HiCCUPS^26^, was applied separately to identify loops at 5 and 10 kb resolutions with default parameters. Finally, loops from two methods were pooled together, which yields a list of 21,738, 12,460 and 16,930 loops in hMSC, adipocytes and osteoblasts, respectively.

### Statistical identification of differential contact loops

To find differential contact loops in adipocytes and osteoblast comparing with hMSC, we first merged loops in chosen cells into a union set by “merge2Dbed.pl” function in HOMER with the default parameters. Next, we counted raw contact frequencies within loops from filtered Hi-C read pairs of each cell replicate, and built a contact frequency matrix with respect to loop sets and replicates. The contact frequency matrix was then used as input in edgeR^54^. After normalizing by the trimmed mean of M values (TMM), differential contact loops between hMSC and adipocytes or osteoblasts were identified using a generalized linear model (GLM) likelihood ratio test. The significance was determined by *P* < 0.01.

### Genomic elements enrichment

Chromatin states from an imputed 25-state model of bone marrow derived MSC (E026), MSC derived adipocyte (E023) and osteoblast (E129) were obtained from the Roadmap Epigenomics project (https://personal.broadinstitute.org/jernst/MODEL_IMPUTED12MARKS/). The annotation details were listed in Supplementary Table 5. We compared the chromatin elements enrichment between Hi-C interacting fragments/loop anchors and other genomic regions, and between interacting fragments/loop anchors and their disjoint 2 kb away regions. For comparing with genomic regions, we focused on enhancer, promoter and positive regulatory associated and additional quiescent annotations (Supplementary Table 4). Firstly, regions were segmented into 200 bp bin pools. Randomly selection was executed 1,000 times, and 1,000 bins were selected from each bin pools at each time. Proportions of bins overlapped with annotation states were calculated at each time for interacting fragments and genomic regions. Z-test was used to find the significant difference of overlapping.

For comparing between the interacting fragments and their disjoint 2 kb away regions, enrichment was estimated by XGR package^55^ implemented in R. 16 annotations associated with transcription, enhancer and promoter were selected to test enrichment in 3 cells (Fig. 2d). Enrichment Z-scores resulting from XGR were plotted to show different enrichment preference between two regions. Statistical significance for comparing CTCF enrichment between loop anchors/interaction fragments and their 2 kb interval regions were calculated with treating 2 kb interval regions as background (Fig. 2c and Supplementary Fig. 3b).

### RNA-seq data processing and differential expression analysis

RNA-seq reads were aligned to human genome (built form the Gencode v19 gene annotation) using STAR^56^ with default parameters. Duplicates were marked by PicardTools (v2.18.9)^57^. Duplicate and low mapping quality reads (MAPQ < 30) were removed for subsequent analyses. The transcript and gene-based expression levels were quantified and normalized to transcript per million (TPM) using RSEM (v1.3.0)^58^. The expected counts of genes/transcripts from RSEM were next normalized by the TMM method. Genes/transcripts that had TMM count >1 in at least 50% of the samples were selected, and were transformed to estimate the mean-variance relationship by voom function implemented in limma package^59^ from R. The data were then tested for differential expression by linear model.

### ChIP-seq data processing and enhancer analysis

ChIP-seq reads were aligned to the human genome assembly (hg19) using Bowtie2^60^ with default settings. Duplicate reads and reads with MAPQ < 30 were discarded. MACS2^61^ was used to perform peak calling with the following parameters (-g hs -p 0.01 --nomodel --extsize 147 --keep-dup all). Peaks of H3K4me1 and H3K27ac were found for each cell replicate separately. Replicated peaks were identified by at least 50% overlap. Putative enhancers were further defined by merging replicated peaks of H3K4me1 and H3K27ac in each cell type.

### Enhancer enrichment in differential contact loops

Overlapped enhancers between adipocytes/osteoblasts and hMSC were defined by at least 80% of the enhancer region in differentiated cell were overlapped with the enhancer in hMSC. The rest were regarded as differential enhancers. BEDtools “intersect” function was used to find overlapped enhancers in adipocytes and osteoblasts. We investigated whether overlapped or differential enhancers were enriched in differential contact loops. Fisher test was applied to find enrichment significance. Enrichment direction was indicated by odds ratio.

### ATAC-seq data processing and peak calling

Adaptors were trimmed from ATAC-seq reads sequences using custom python scripts. Pair-end reads were aligned to hg19 using Bowtie2. Duplicate reads and reads with MAPQ < 30 were discarded. Reads mapping to the mitochondria and chromosome Y were removed. After filtering, the qualified reads were subjected to MACS2 to call peaks for each sample with parameters (-q 0.05 --nomodel --shift -100 --extsize 200 --keep-dup all). Peaks mapped to the consensus excludable ENCODE blacklist (http://hgdownload.cse.ucsc.edu/goldenPath/hg19/encodeDCC/wgEncodeMapability/) were filtered. The peaks between replicates of the same cell type were merged using BEDTools^62^. In total, we identified 138,820 and 120,209 peaks from adipocytes and osteoblasts, respectively. In order to compare TFs footprints of adipocytes and osteoblasts with hMSC and another unrelative cell, we obtained ATAC-seq peaks information of hMSC from Rauch et al. (GSE113253)^15^ and GM12878 cell line from Buenrostro *et al.* (GSE47753)^51^.

### Colocalization between ATAC-seq and H3K27ac ChIP-seq

Complementary genomic regions to ATAC-seq peaks were selected for adipocytes and osteoblasts, from which peak length matching regions were randomly generated. The GC contents of random regions were calculated by BEDtools. Regions with GC content matching with peaks were integrated to construct the matching region set. ATAC-seq reads and H3K27ac ChIP-seq reads mapped to ATAC peaks or matching regions were counted and normalized by RPKM in each cell using deepTools software^63^. Colocalization profiles were plotted at a 10 kb region flanking the ATAC peak summits/region midpoints.

### TF motifs enrichment in ATAC-seq peaks

The HOMER motif finding function “findMotifsGenome.pl” was used to detect enriched TF motifs in ATAC-seq peaks with parameters (-size 200 -mask) and the hg19 genome reference. For background chosen, we found motifs within ATAC-seq peaks identified for hMSC, adipocytes, osteoblasts and GM12878 using the union peak set as background. 413 known motifs available in HOMER were used to test for enrichment. The enrichment Z-scores were used to compare and find cell-regulatory motifs across cells.

### Regulatory networks construction

We constructed regulatory networks using multi-omic data, including loop structures, gene expression levels, enhancers and chromatin accessible regions, as well as TFs binding sites collecting by Yevshin *et al.*^45^. The anchors were firstly rescaled to 10 kb, and then searched for ATAC-seq peaks. Unique loops with both anchors mapping with ATAC-seq peaks were kept. The unique gene promoters were consisted of −2 kb to +1 kb regions to TSS of each gene transcript. Next, one side of loop anchors was mapped with those promoters while the other side was mapped with unique enhancers. Both promoters and enhancers were then mapped with TFs binding sites. By this way, the gene and TFs were connected, and the edge weight was defined as:

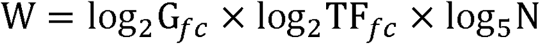

Where G_fc_ and TF_fc_ refer to expression fold change of target gene and TFs after differentiation. N refers to the sum of peak caller numbers that are able to recognize the binding events at promoters and enhancers. The node weight was defined as the expression fold change. The TFs binding sites and target genes were next utilized to search for eQTLs (*P* < 0.05). The eQTLs data from subcutaneous and visceral omentum adipose, LCLs and whole blood were derived from GTEx database (v8)^64^. SNPs located at the TFs binding sites and effecting the same genes as the loop anchors interacting with were subsequentially added to the networks. The weights between SNPs and TFs were defined as −log10 transformed eQTL *P* values.

## Acknowledgements

This work was supported by grants from the National Natural Science Foundation of China (31871264, 31970569), the Natural Science Basic Research Program Shaanxi Province (2019JM-119) and the Fundamental Research Funds for the Central Universities. We would like to thank the GTEx Consortium. We obtained GTEx data through dbGaP authorized access at https://dbgap.ncbi.nlm.nih.gov/aa/wga.cgi?page=login with the accession number of phs000424.v8.p2.

## Author contributions

T.L.Y. and Y.G. conceived and supervised this project. R.H.H. conducted the computational work. J.G. and R.H.H. performed the cell culture experiments. Y.R. performed visualization. S.S.D. built the pipeline for Hi-C data analysis. H.C. and D.L.Z. carried out the library construction experiments. Y.X.C. participated in data analysis. R.H.H. wrote the manuscript with the assistance of other authors.

## Conflict of Interest

All the authors declare that they have no conflicts of interest.

## Notes

### Competing Interest Statement

The authors have declared no competing interest.

## References

1. Brunmeir R, Wu J, Peng X, Kim SY, Julien SG, Zhang Q, et al. Comparative Transcriptomic and Epigenomic Analyses Reveal New Regulators of Murine Brown Adipogenesis. PLoS genetics 2016, 12(12): e1006474.

2. Menssen A, Haupl T, Sittinger M, Delorme B, Charbord P, Ringe J. Differential gene expression profiling of human bone marrow-derived mesenchymal stem cells during adipogenic development. BMC genomics 2011, 12: 461.

3. Zhang W, Dong R, Diao S, Du J, Fan Z, Wang F. Differential long noncoding RNA/mRNA expression profiling and functional network analysis during osteogenic differentiation of human bone marrow mesenchymal stem cells. Stem cell research & therapy 2017, 8(1): 30.

4. Piek E, Sleumer LS, van Someren EP, Heuver L, de Haan JR, de Grijs I, et al. Osteo-transcriptomics of human mesenchymal stem cells: accelerated gene expression and osteoblast differentiation induced by vitamin D reveals c-MYC as an enhancer of BMP2-induced osteogenesis. Bone 2010, 46(3): 613–627.

5. Scheideler M, Elabd C, Zaragosi LE, Chiellini C, Hackl H, Sanchez-Cabo F, et al. Comparative transcriptomics of human multipotent stem cells during adipogenesis and osteoblastogenesis. BMC genomics 2008, 9: 340.

6. Siersbaek R, Nielsen R, Mandrup S. PPARgamma in adipocyte differentiation and metabolism-- novel insights from genome-wide studies. FEBS letters 2010, 584(15): 3242–3249.

7. Lehrke M, Lazar MA. The many faces of PPARgamma. Cell 2005, 123(6): 993–999.

8. Lefterova MI, Haakonsson AK, Lazar MA, Mandrup S. PPARgamma and the global map of adipogenesis and beyond. Trends in endocrinology and metabolism: TEM 2014, 25(6): 293–302.

9. Rosen ED, Hsu CH, Wang X, Sakai S, Freeman MW, Gonzalez FJ, et al. C/EBPalpha induces adipogenesis through PPARgamma: a unified pathway. Genes & development 2002, 16(1): 22–26.

10. Madsen MS, Siersbaek R, Boergesen M, Nielsen R, Mandrup S. Peroxisome proliferator-activated receptor gamma and C/EBPalpha synergistically activate key metabolic adipocyte genes by assisted loading. Molecular and cellular biology 2014, 34(6): 939–954.

11. Ducy P, Zhang R, Geoffroy V, Ridall AL, Karsenty G. Osf2/Cbfa1: a transcriptional activator of osteoblast differentiation. Cell 1997, 89(5): 747–754.

12. Nakashima K, Zhou X, Kunkel G, Zhang Z, Deng JM, Behringer RR, et al. The novel zinc finger-containing transcription factor osterix is required for osteoblast differentiation and bone formation. Cell 2002, 108(1): 17–29.

13. Meyer MB, Benkusky NA, Sen B, Rubin J, Pike JW. Epigenetic Plasticity Drives Adipogenic and Osteogenic Differentiation of Marrow-derived Mesenchymal Stem Cells. The Journal of biological chemistry 2016, 291(34): 17829–17847.

14. Wang L, Niu N, Li L, Shao R, Ouyang H, Zou W. H3K36 trimethylation mediated by SETD2 regulates the fate of bone marrow mesenchymal stem cells. PLoS biology 2018, 16(11): e2006522.

15. Rauch A, Haakonsson AK, Madsen JGS, Larsen M, Forss I, Madsen MR, et al. Osteogenesis depends on commissioning of a network of stem cell transcription factors that act as repressors of adipogenesis. Nature genetics 2019, 51(4): 716–727.

16. Ke Y, Xu Y, Chen X, Feng S, Liu Z, Sun Y, et al. 3D Chromatin Structures of Mature Gametes and Structural Reprogramming during Mammalian Embryogenesis. Cell 2017, 170(2): 367–381 e320.

17. Dixon JR, Selvaraj S, Yue F, Kim A, Li Y, Shen Y, et al. Topological domains in mammalian genomes identified by analysis of chromatin interactions. Nature 2012, 485(7398): 376–380.

18. Sun JH, Zhou L, Emerson DJ, Phyo SA, Titus KR, Gong W, et al. Disease-Associated Short Tandem Repeats Co-localize with Chromatin Domain Boundaries. Cell 2018, 175(1): 224–238 e215.

19. Sexton T, Cavalli G. The role of chromosome domains in shaping the functional genome. Cell 2015, 160(6): 1049–1059.

20. Greenwald WW, Chiou J, Yan J, Qiu Y, Dai N, Wang A, et al. Pancreatic islet chromatin accessibility and conformation reveals distal enhancer networks of type 2 diabetes risk. Nature communications 2019, 10(1): 2078.

21. Nolis IK, McKay DJ, Mantouvalou E, Lomvardas S, Merika M, Thanos D. Transcription factors mediate long-range enhancer-promoter interactions. Proceedings of the National Academy of Sciences of the United States of America 2009, 106(48): 20222–20227.

22. Jin F, Li Y, Dixon JR, Selvaraj S, Ye Z, Lee AY, et al. A high-resolution map of the three-dimensional chromatin interactome in human cells. Nature 2013, 503(7475): 290–294.

23. Greenwald WW, Li H, Benaglio P, Jakubosky D, Matsui H, Schmitt A, et al. Subtle changes in chromatin loop contact propensity are associated with differential gene regulation and expression. Nature communications 2019, 10(1): 1054.

24. Wingett S, Ewels P, Furlan-Magaril M, Nagano T, Schoenfelder S, Fraser P, et al. HiCUP: pipeline for mapping and processing Hi-C data. F1000Research 2015, 4: 1310.

25. Lieberman-Aiden E, van Berkum NL, Williams L, Imakaev M, Ragoczy T, Telling A, et al. Comprehensive mapping of long-range interactions reveals folding principles of the human genome. Science 2009, 326(5950): 289–293.

26. Rao SS, Huntley MH, Durand NC, Stamenova EK, Bochkov ID, Robinson JT, et al. A 3D map of the human genome at kilobase resolution reveals principles of chromatin looping. Cell 2014, 159(7): 1665–1680.

27. Bult CJ, Blake JA, Smith CL, Kadin JA, Richardson JE, Mouse Genome Database G. Mouse Genome Database (MGD) 2019. Nucleic acids research 2019, 47(D1): D801–D806.

28. Smallwood A, Ren B. Genome organization and long-range regulation of gene expression by enhancers. Current opinion in cell biology 2013, 25(3): 387–394.

29. Siersbaek R, Madsen JGS, Javierre BM, Nielsen R, Bagge EK, Cairns J, et al. Dynamic Rewiring of Promoter-Anchored Chromatin Loops during Adipocyte Differentiation. Molecular cell 2017, 66(3): 420–435 e425.

30. Rubin AJ, Barajas BC, Furlan-Magaril M, Lopez-Pajares V, Mumbach MR, Howard I, et al. Lineage-specific dynamic and pre-established enhancer-promoter contacts cooperate in terminal differentiation. Nature genetics 2017, 49(10): 1522–1528.

31. Nora EP, Lajoie BR, Schulz EG, Giorgetti L, Okamoto I, Servant N, et al. Spatial partitioning of the regulatory landscape of the X-inactivation centre. Nature 2012, 485(7398): 381–385.

32. Morris JA, Tsai PC, Joehanes R, Zheng J, Trajanoska K, Soerensen M, et al. Epigenome-wide Association of DNA Methylation in Whole Blood With Bone Mineral Density. Journal of bone and mineral research : the official journal of the American Society for Bone and Mineral Research 2017, 32(8): 1644–1650.

33. Greenblatt MB, Shim JH. Osteoimmunology: a brief introduction. Immune network 2013, 13(4): 111–115.

34. Fulzele K, Riddle RC, DiGirolamo DJ, Cao X, Wan C, Chen D, et al. Insulin receptor signaling in osteoblasts regulates postnatal bone acquisition and body composition. Cell 2010, 142(2): 309–319.

35. Ohta H, Itoh N. Roles of FGFs as Adipokines in Adipose Tissue Development, Remodeling, and Metabolism. Frontiers in endocrinology 2014, 5: 18.

36. Ott CJ, Federation AJ, Schwartz LS, Kasar S, Klitgaard JL, Lenci R, et al. Enhancer Architecture and Essential Core Regulatory Circuitry of Chronic Lymphocytic Leukemia. Cancer cell 2018, 34(6): 982–995 e987.

37. Consortium EP. An integrated encyclopedia of DNA elements in the human genome. Nature 2012, 489(7414): 57–74.

38. Cadoudal T, Distel E, Durant S, Fouque F, Blouin JM, Collinet M, et al. Pyruvate dehydrogenase kinase 4: regulation by thiazolidinediones and implication in glyceroneogenesis in adipose tissue. Diabetes 2008, 57(9): 2272–2279.

39. Miki H, Yamauchi T, Suzuki R, Komeda K, Tsuchida A, Kubota N, et al. Essential role of insulin receptor substrate 1 (IRS-1) and IRS-2 in adipocyte differentiation. Molecular and cellular biology 2001, 21(7): 2521–2532.

40. Wang P, Keijer J, Bunschoten A, Bouwman F, Renes J, Mariman E. Insulin modulates the secretion of proteins from mature 3T3-L1 adipocytes: a role for transcriptional regulation of processing. Diabetologia 2006, 49(10): 2453–2462.

41. Attane C, Esteve D, Chaoui K, Iacovoni JS, Corre J, Moutahir M, et al. Human Bone Marrow Is Comprised of Adipocytes with Specific Lipid Metabolism. Cell reports 2020, 30(4): 949–958 e946.

42. Komori T. Molecular Mechanism of Runx2-Dependent Bone Development. Molecules and cells 2020.

43. Hiruma Y, Tsuda E, Maeda N, Okada A, Kabasawa N, Miyamoto M, et al. Impaired osteoclast differentiation and function and mild osteopetrosis development in Siglec-15-deficient mice. Bone 2013, 53(1): 87–93.

44. Pei YF, Liu L, Liu TL, Yang XL, Zhang H, Wei XT, et al. Joint Association Analysis Identified 18 New Loci for Bone Mineral Density. Journal of bone and mineral research : the official journal of the American Society for Bone and Mineral Research 2019, 34(6): 1086–1094.

45. Yevshin I, Sharipov R, Kolmykov S, Kondrakhin Y, Kolpakov F. GTRD: a database on gene transcription regulation-2019 update. Nucleic acids research 2019, 47(D1): D100–D105.

46. Khalid AB, Krum SA. Estrogen receptors alpha and beta in bone. Bone 2016, 87: 130–135.

47. Watters RJ, Hartmaier RJ, Osmanbeyoglu HU, Gillihan RM, Rae JM, Liao L, et al. Steroid receptor coactivator-1 can regulate osteoblastogenesis independently of estrogen. Molecular and cellular endocrinology 2017, 448: 21–27.

48. Schneider CA, Rasband WS, Eliceiri KW. NIH Image to ImageJ: 25 years of image analysis. Nature methods 2012, 9(7): 671–675.

49. Gayral P, Weinert L, Chiari Y, Tsagkogeorga G, Ballenghien M, Galtier N. Next-generation sequencing of transcriptomes: a guide to RNA isolation in nonmodel animals. Molecular ecology resources 2011, 11(4): 650–661.

50. Chen XF, Zhu DL, Yang M, Hu WX, Duan YY, Lu BJ, et al. An Osteoporosis Risk SNP at 1p36.12 Acts as an Allele-Specific Enhancer to Modulate LINC00339 Expression via Long-Range Loop Formation. American journal of human genetics 2018, 102(5): 776–793.

51. Buenrostro JD, Giresi PG, Zaba LC, Chang HY, Greenleaf WJ. Transposition of native chromatin for fast and sensitive epigenomic profiling of open chromatin, DNA-binding proteins and nucleosome position. Nature methods 2013, 10(12): 1213–1218.

52. Imakaev M, Fudenberg G, McCord RP, Naumova N, Goloborodko A, Lajoie BR, et al. Iterative correction of Hi-C data reveals hallmarks of chromosome organization. Nature methods 2012, 9(10): 999–1003.

53. Heinz S, Benner C, Spann N, Bertolino E, Lin YC, Laslo P, et al. Simple combinations of lineage-determining transcription factors prime cis-regulatory elements required for macrophage and B cell identities. Molecular cell 2010, 38(4): 576–589.

54. Robinson MD, McCarthy DJ, Smyth GK. edgeR: a Bioconductor package for differential expression analysis of digital gene expression data. Bioinformatics 2010, 26(1): 139–140.

55. Fang H, Knezevic B, Burnham KL, Knight JC. XGR software for enhanced interpretation of genomic summary data, illustrated by application to immunological traits. Genome medicine 2016, 8(1): 129.

56. Dobin A, Davis CA, Schlesinger F, Drenkow J, Zaleski C, Jha S, et al. STAR: ultrafast universal RNA-seq aligner. Bioinformatics 2013, 29(1): 15–21.

57. DePristo MA, Banks E, Poplin R, Garimella KV, Maguire JR, Hartl C, et al. A framework for variation discovery and genotyping using next-generation DNA sequencing data. Nature genetics 2011, 43(5): 491–498.

58. Li B, Dewey CN. RSEM: accurate transcript quantification from RNA-Seq data with or without a reference genome. BMC bioinformatics 2011, 12: 323.

59. Ritchie ME, Phipson B, Wu D, Hu Y, Law CW, Shi W, et al. limma powers differential expression analyses for RNA-sequencing and microarray studies. Nucleic acids research 2015, 43(7): e47.

60. Langmead B, Salzberg SL. Fast gapped-read alignment with Bowtie 2. Nature methods 2012, 9(4): 357–359.

61. Zhang Y, Liu T, Meyer CA, Eeckhoute J, Johnson DS, Bernstein BE, et al. Model-based analysis of ChIP-Seq (MACS). Genome biology 2008, 9(9): R137.

62. Quinlan AR, Hall IM. BEDTools: a flexible suite of utilities for comparing genomic features. Bioinformatics 2010, 26(6): 841–842.

63. Ramirez F, Ryan DP, Gruning B, Bhardwaj V, Kilpert F, Richter AS, et al. deepTools2: a next generation web server for deep-sequencing data analysis. Nucleic acids research 2016, 44(W1): W160–165.

64. Consortium GT. Human genomics. The Genotype-Tissue Expression (GTEx) pilot analysis: multitissue gene regulation in humans. Science 2015, 348(6235): 648–660.

