## Supplemental Figures for "3D chromatin organization changes modulate adipogenesis and osteogenesis"

Supplementary Figures

Ruo-Han Hao^1^, Yan Guo^1^, Jing Guo^1^, Yu Rong^1^, Shi Yao^1^, Yi-Xiao Chen^1^, Shan-Shan Dong^1^, Dong-Li Zhu^1^, Hao Chen^1^, Tie-Lin Yang^1^*

^1^Key Laboratory of Biomedical Information Engineering of Ministry of Education, Biomedical Informatics & Genomics Center, School of Life Science and Technology, Xi'an Jiaotong University, Xi'an, Shaanxi, P. R. China, 710049

*Corresponding authors: Tie-Lin Yang, Ph.D.

Key Laboratory of Biomedical Information Engineering of Ministry of Education, Biomedical Informatics & Genomics Center, School of Life Science and Technology, Xi'an Jiaotong University, Xi'an, Shaanxi, P. R. China, 710049

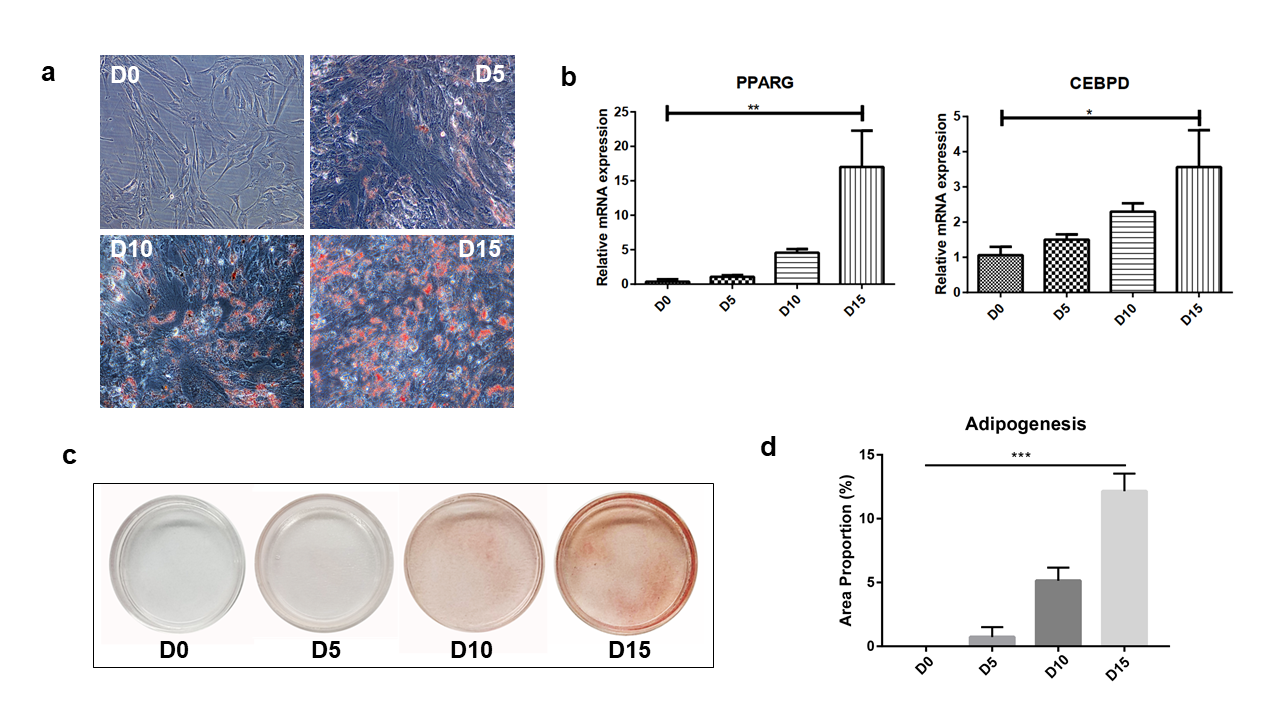

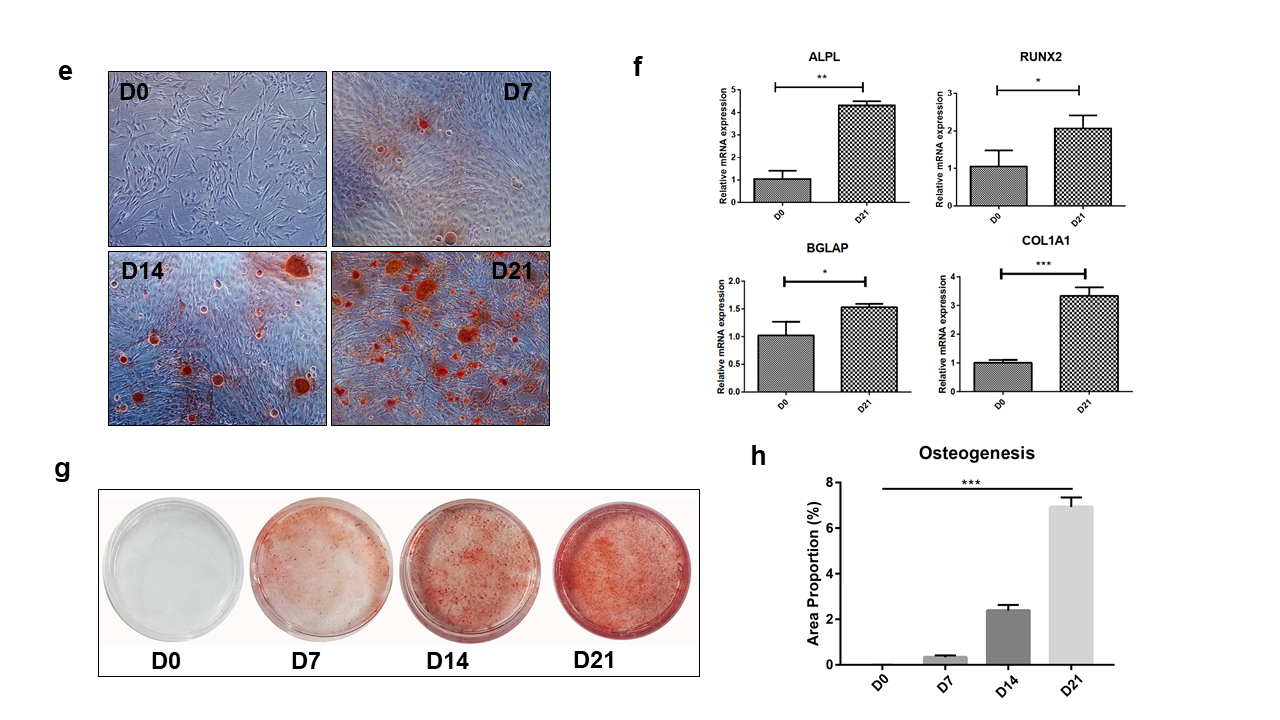

Supplementary Fig. 1. hMSC differentiation status verification by microscopic examination, qRT-PCR of marker genes and cell staining. **a-d** Adipogenesis status were checked at time points day 0, 5, 10 and 15; **e-h** osteogenesis status were checked at time points day 0, 7, 14 and 21. Staining area proportion in **d** and **h** was calculated by ImageJ software from images taken under microscope. The statistical significance was indicated by t-test.

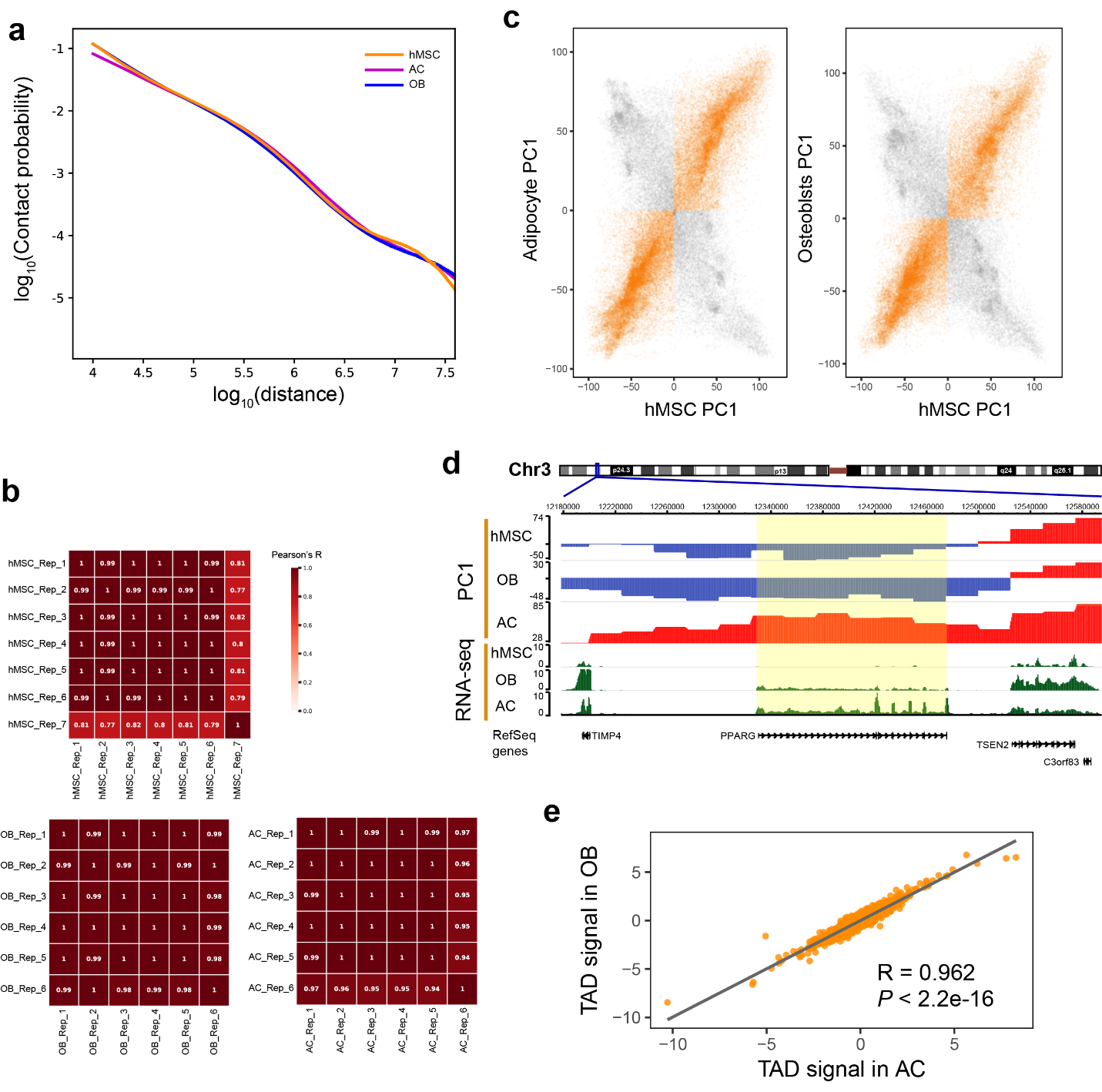

Supplementary Fig. 2. **a** The average contact probability decreases with the genomic distance increase. **b** Heatmaps showing the correlation coefficients of Hi-C replicates for each cell. **c** Chromatin compartment consistency and switch after hMSC differentiation. **d** The genome browser screenshot showing the compartment A/B states and RNA-seq signals surrounding *PPARG* gene between 3 cells. **e** The correlation of TAD signals between AC and OB. *P* values and correlation coefficient displayed here were estimated by Pearson correlation test.

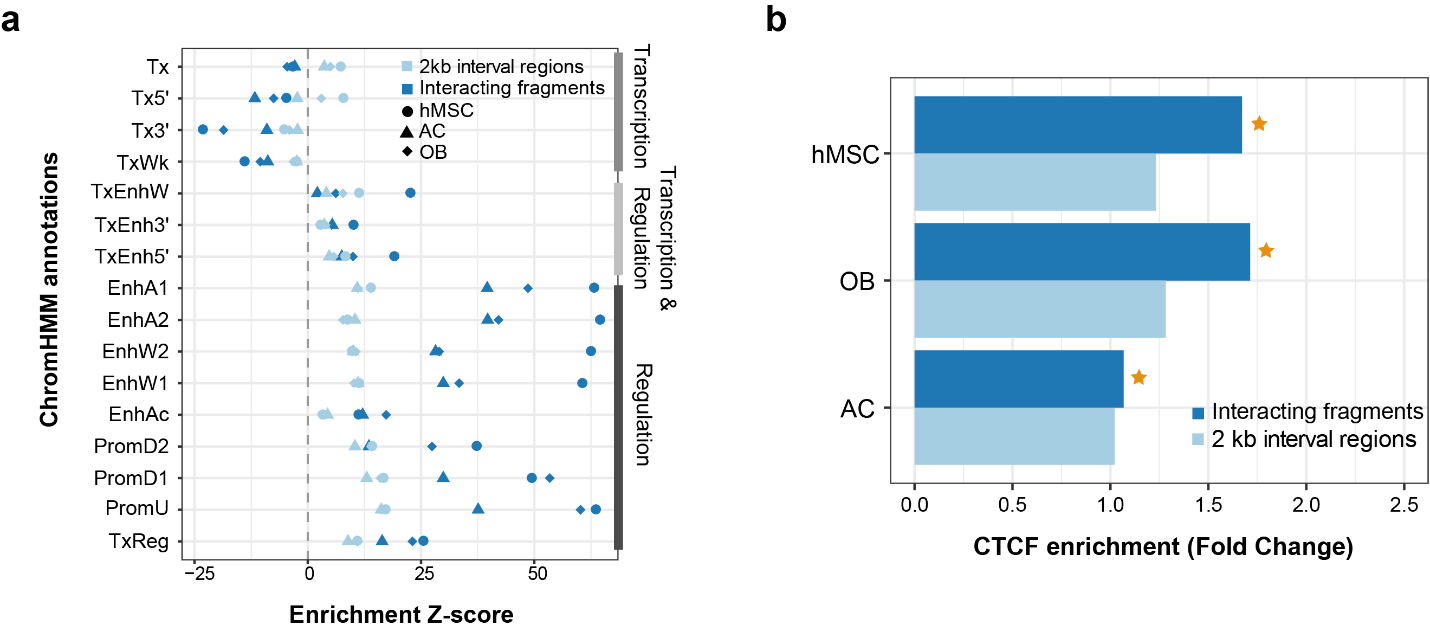

Supplementary Fig. 3. **a** Region enrichment results illustrating the ChromHMM annotation enrichment at interacting fragments and their 2 kb interval regions. Enrichment Z-scores are plotted. Cells are separated by different shapes, and regions are distinguished by different colors. **b** Bar plot showing the fold change of CTCF binding sites enrichment at interacting fragments and their 2 kb interval regions. Statistical significance was calculated with treating 2 kb interval regions as background. *P* < 0.05 is asterisked.

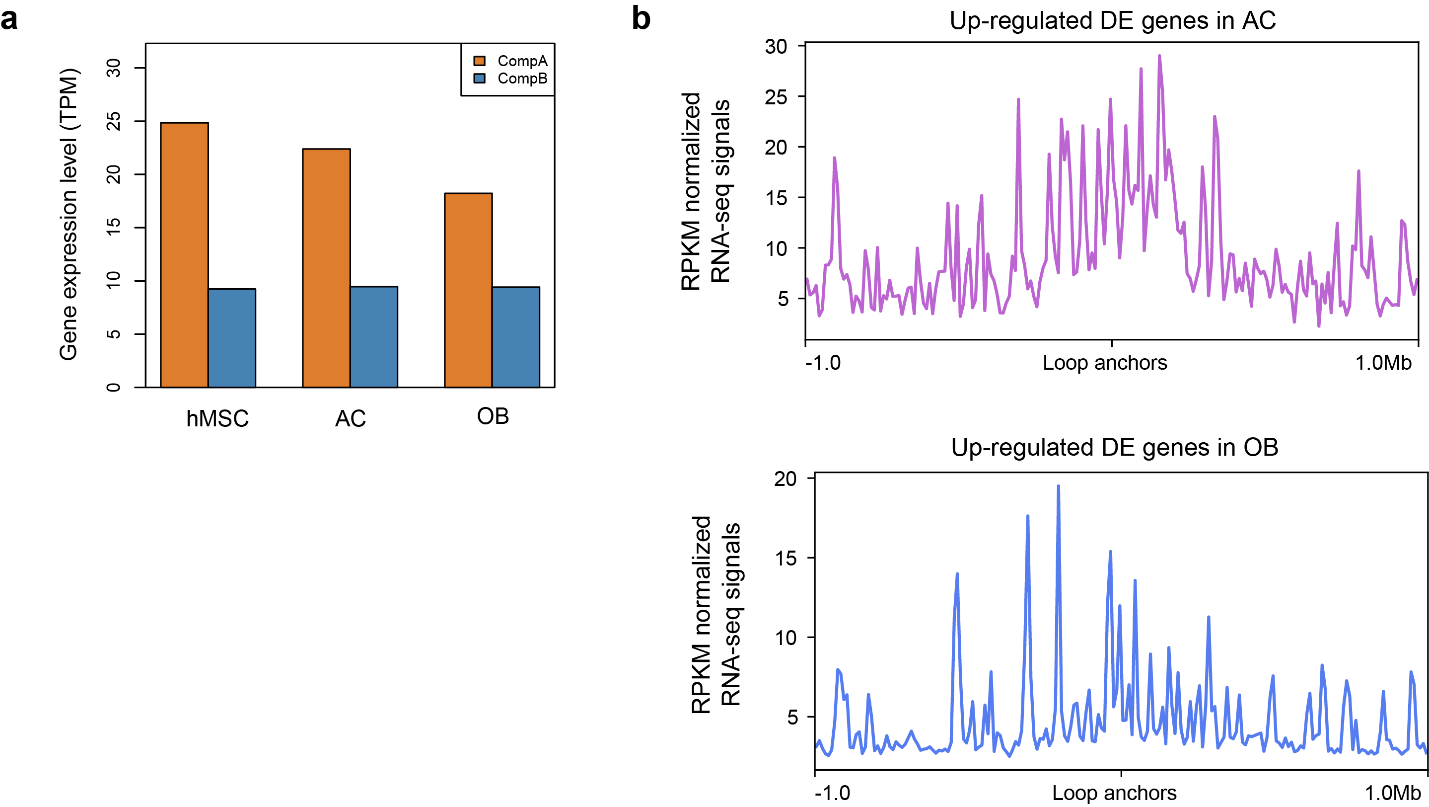

Supplementary Fig. 4. **a** The comparison of average gene expression levels (normalized using TPM) between A and B compartments. **b** The transcription signal distribution surrounding loop anchors in AC and OB.

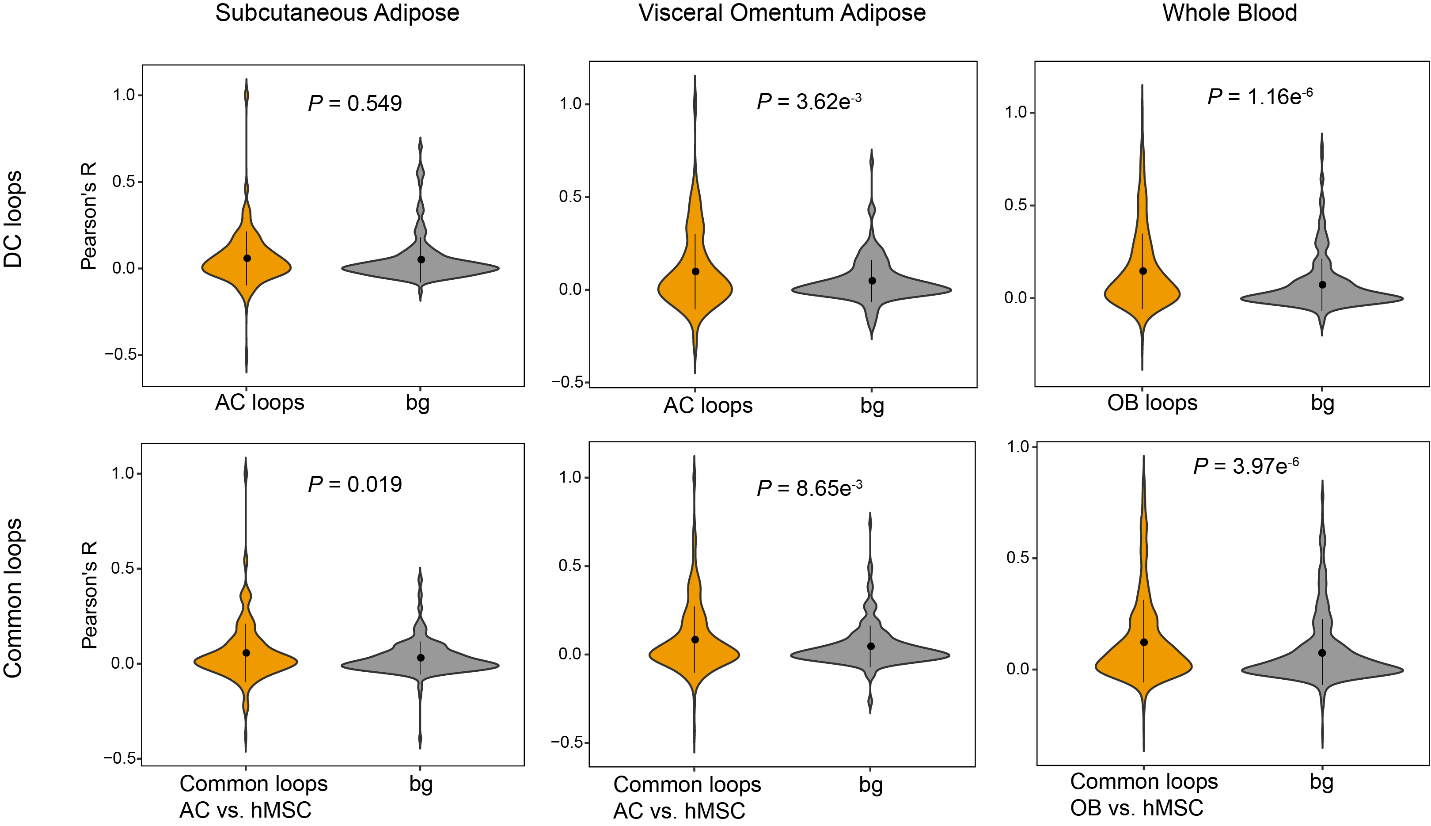

Supplementary Fig. 5. Violin plots showing the distribution of gene co-expression coefficients within loops and background groups. *P* values were indicated by t-test.

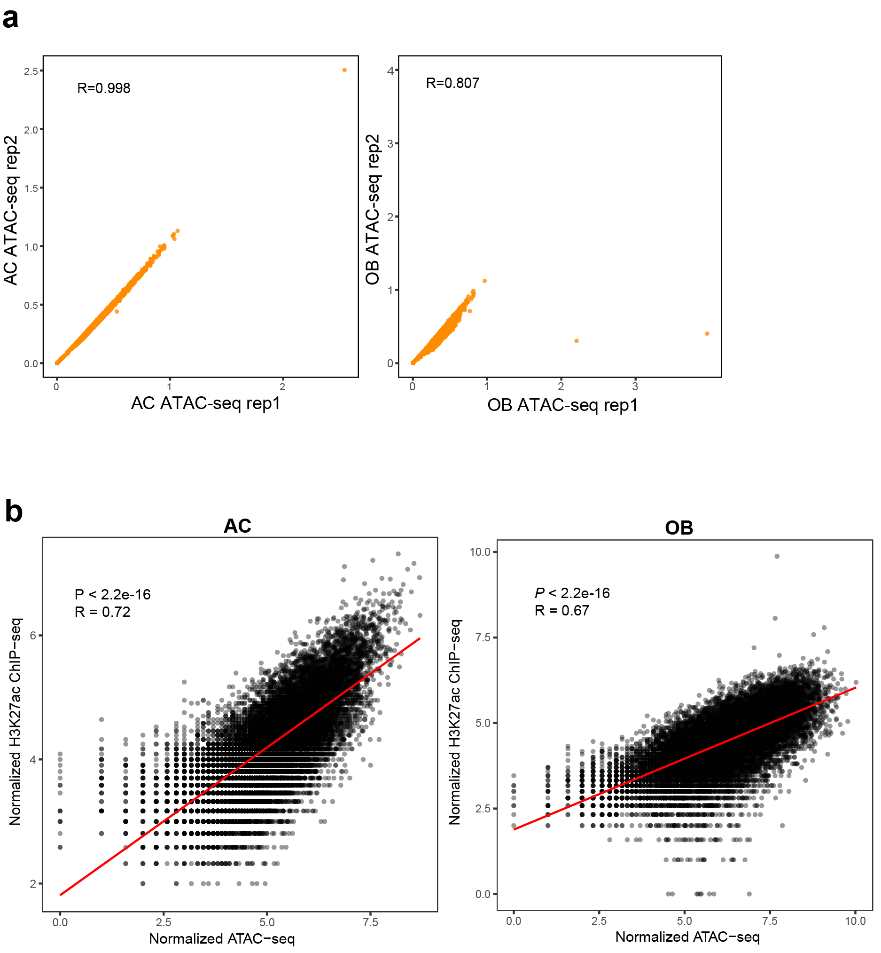

Supplementary Fig. 6. **a** The correlation between ATAC-seq replicates for AC and OB. “R” indicates Pearson correlation coefficient. **b** The correlation between ATAC-seq and H3K27ac ChIP-seq. Pearson correlation test was executed to evaluate correlation significance.

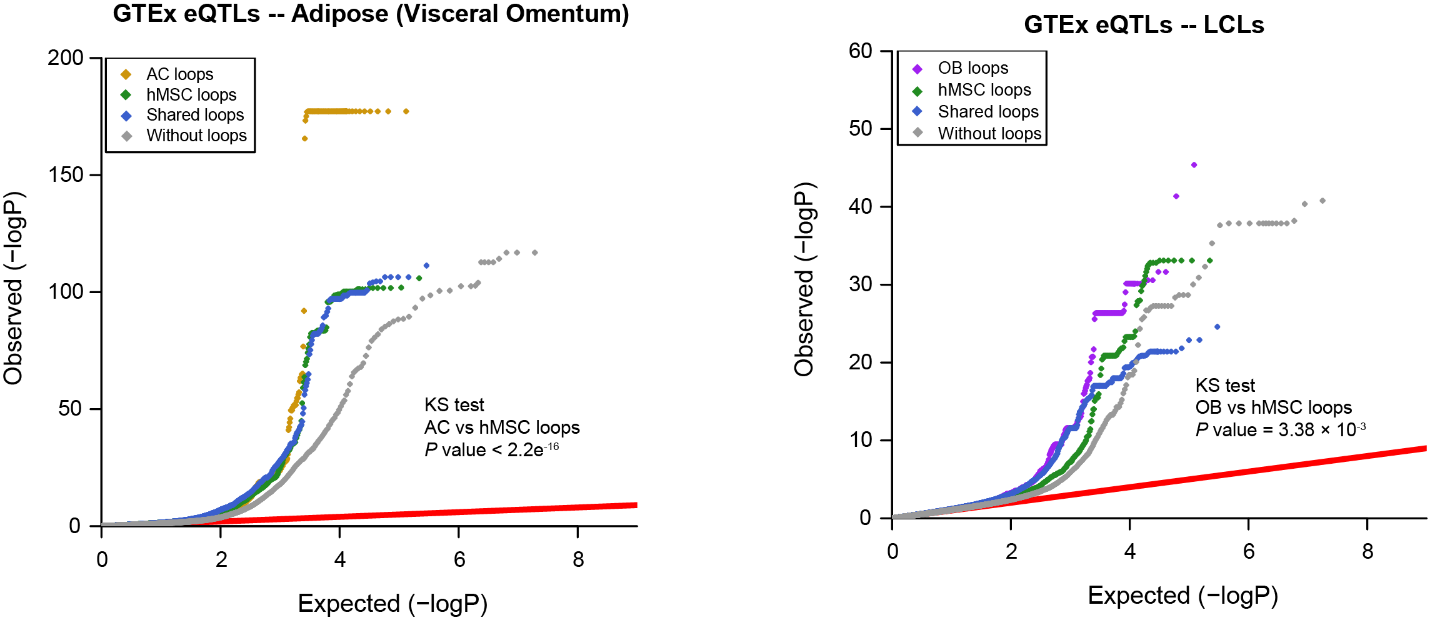

Supplementary Fig. 7. Q-Q plots drawn with eQTL data from visceral omentum adipose (left) and LCLs (right) illustrating the significant enrichment of eQTL associations at AC/OB loops.
